# Aging-associated vacuolation of multi-ciliated cells in the distal mouse oviduct reflects unique cell identity and luminal microenvironment

**DOI:** 10.1101/2024.09.07.611808

**Authors:** Keerthana Harwalkar, Nobuko Yamanaka, Alain S Pacis, Selina Zhao, Katie Teng, Warwick Pitman, Mitaali Taskar, Vera Lynn, Alex Frances Thornton, Matthew J Ford, Yojiro Yamanaka

## Abstract

The female reproductive organs present with the earliest aging characteristics, such as a decline in fertility and estrous cyclicity. While age-related changes in the ovary are well-documented, it is unclear if any age-associated changes occur in the other female reproductive organs, such as the oviduct/fallopian tube. The recent recognition of the distal end of the fallopian tube as the tissue of origin of high-grade serous tubal ovarian carcinomas (HGSCs), and that its patient demographic is strongly biased to postmenopausal women, motivated us to investigate age-associated changes in this organ.

At the distal end of aged oviducts in mice, we found vacuolated multi-ciliated cells (MCCs) with a severely apically displaced and deformed nucleus. This phenotype was unique to the distal oviduct epithelium –infundibulum (INF) and ampulla (AMP). Ovariectomy did not affect the timeline of MCC vacuolation, suggesting little involvement of ovulation and hormonal regulation. MCC vacuolation was induced in hypoxia or hydroxyurea treatments in *in vitro* organotypic culture of all oviduct regions, not limited to the INF/AMP epithelium. This suggests high oxygen demand in MCCs, compared to other cell types, and a uniquely stressed INF/AMP epithelial microenvironment *in vivo*. We found that the blood circulation of INF/AMP depended on the ovarian artery, different from the rest of the oviduct epithelium and its circulation declined along with ovarian activities. We conclude that a decline in local blood circulation and distinct cellular identity of the INF/AMP epithelium caused age-associated MCC vacuolation, reflecting its mild, chronically stressed microenvironment.

## INTRODUCTION

The oviduct is the tube that extends from the uterus to the ovary, with 4 regions demarcated by their mucosal fold morphologies, epithelial cell transcription factor expression, and proportion/distribution of multi-ciliated cells (MCCs). These regions are the infundibulum/ampulla (INF/AMP), ampulla-isthmus junction (AIJ), isthmus (ISM), and uterotubal junction (UTJ) from distal to proximal, relative to the uterus (1, 2). Gamete and preimplantation embryo movement occurs in the oviduct, aided by the mucosal epithelium (3, 4) and smooth muscle contractions (5). These functions are controlled by hormonal regulation, via the hypothalamic-pituitary-ovary axis, in a cyclic manner over the course of 4-5 days, called the estrous cycle in mice. Peak estrous cyclicity is noted at around 5-months of age in single housed, unmated, inbred C57BL/6J mice, and a decline in this cyclicity occurs from ∼9 months. The initial phase of decline that occurs at 10-12-months of age is characterized by a progressive lengthening of estrous cycles (6). Alongside this loss of estrous cyclicity, a decline in oocyte quality, impaired decidualization, uterine fibrosis (7, 8), prolongation of gestation/labour, and decreased litter size (9) are noted.

Although classified as a type of ovarian cancer, High Grade Serous Ovarian Carcinomas (HGSCs) originate from the distal fimbrial region of the fallopian tubes (10, 11). The frequency of ovulation across the reproductive lifespan is considered as a risk factor because of repetitive exposures to inflammatory cytokines and reactive oxygen species (ROS), that can induce mutations in the adjacent fimbrial epithelium (12). On the other hand, the distal INF/AMP epithelium is developmentally and transcriptionally distinct from the rest of the oviduct epithelia (1, 13). Therefore, this raises a possibility that the INF/AMP epithelial cells are uniquely susceptible to malignant transformation, a concept known as cellular pliancy (14). HGSCs are diagnosed later in life, in postmenopausal individuals typically over 50 years of age (15), suggesting that aging may provide a unique permissive environment for HGSC initiation and/or progression.

In this study, we identified MCC-specific vacuolation in the INF/AMP epithelium of mice at around 7 months of age. Size and frequency of these MCC vacuoles increased with age. Ovariectomy did not change the timeline of vacuolation, suggesting no association with repetitive ovulation or hormonal regulation. Hypoxia and hydroxyurea (HU) treatments in *in vitro* organotypic cultures induced vacuolation in MCCs of all oviduct regions, not limited to the INF/AMP epithelium. This confirms the general high energy demand of MCCs but simultaneously suggests a unique INF/AMP microenvironment *in vivo*. We realized that the INF/AMP blood circulation was coupled to that of the ovary via the ovarian artery and demonstrated that blood circulation into the ovary and INF/AMP was reduced in aged females. Thus, we conclude that MCC vacuolation in the INF/AMP epithelium of aged females is primarily due to constant high-energy demand of MCCs compared to other cell types. MCC vacuolation unique to the INF/AMP epithelium was caused by its distinct cellular property and a mild, chronically stressed INF/AMP microenvironment, due to unique circulation patterns. We propose that this could contribute to age-associated vulnerability to HGSC.

## RESULTS

### Epithelium-lined diverticula along the aged mouse oviduct

To investigate any age-related structural changes in the mouse oviduct, we examined whole-mount and 3D confocal images of oviducts isolated from 7-month-old (7mo), 13-month-old (13mo), and 18-month-old (18mo) females. 7mo oviducts did not present with any obvious structural changes (Fig. 1A, B), relative to 2-4mo oviducts (1). Outpouching of the smooth muscle layer was noted in 13mo oviducts, predominantly in the proximal AIJ and ISM regions (Fig. 1C, E), but not evident in the distal INF/AMP (Fig. 1D). In oviducts isolated from 18mo mice, the distal INF/AMP region presented with outpouching of the smooth muscle (Fig. 1G), similar to those noted in the proximal oviducts of 13mo mice (Fig. 1E). Large outpouched cysts were restricted to the proximal oviduct (Fig. 1F, H). Interestingly, these cysts observed throughout the tube in 18mo mice were lined with E-CAD+ve epithelium that connected into the oviduct lumen (Fig. 1I-K), indicating that these structures are diverticula. Indeed, the outpouched epithelium that faced the lumen was SOX17 positive (Fig. 1I-K), confirming that it originated from the oviduct epithelium (1). It is likely caused by tears in the smooth muscle layer, potentially due to the stiffened tube structure (16) and estrous cycle-associated changes in luminal pressure. Age associated fibrosis, muscle layer thickness, and continued peristaltic movements of the oviduct could contribute to the formation of diverticula; in particular, the large diverticula observed in the ISM and UTJ.

**Figure 1:**
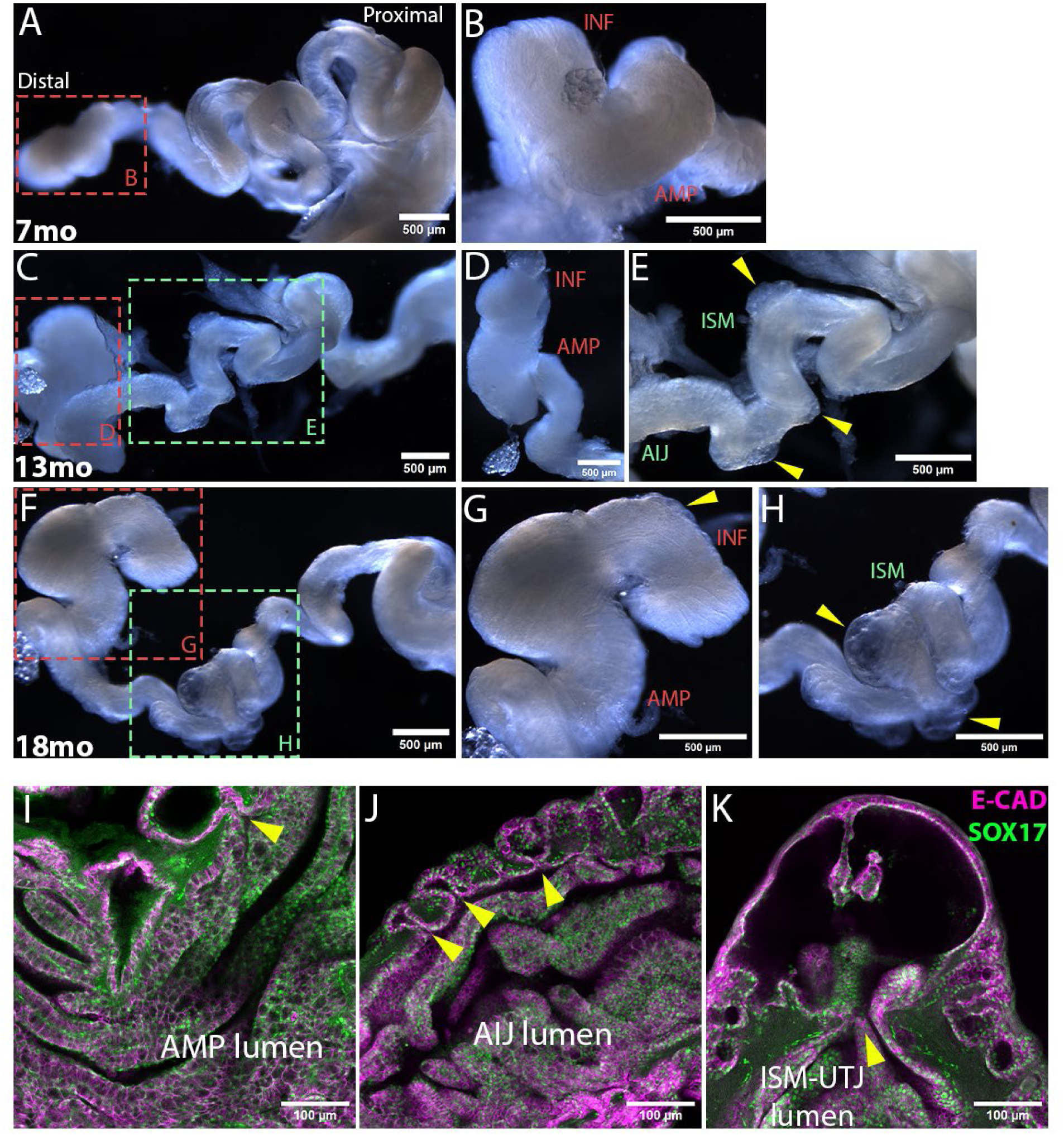
Epithelial outpouching along the aged mouse oviduct. (A-H) Whole-mount images of an uncoiled oviduct dissected from a 7mo (A, B), 13mo (B-D) and 18mo (E-H) mouse showing outpouching (yellow arrows in E, G, H). Aged 13mo and 18mo oviducts were stiffer and less translucent than 7mo oviducts. (I-K) 3D confocal images of an 18mo AMP (I), AIJ (J), and ISM-UTJ (K) regions showing E-CAD and SOX17-expressing epithelial cells lined the diverticula and connected into the oviduct lumen (yellow arrows in I-K).

### Large cytoplasmic vacuole with an apically displaced, crescent-shaped deformed nucleus was uniquely noted in INF/AMP MCCs of the aged oviduct

To investigate any age-related changes at the cellular level, we examined transverse sections of oviducts isolated from 4-month-old (4mo) and 18mo females from four distinct regions: INF/AMP, AIJ, ISM and UTJ (Fig. 2A-D, K-P). In the 4mo INF/AMP epithelium, the nucleus of epithelial cells was round or oval shaped and positioned basally (Fig. 2A, B). Many epithelial cells in the 18mo INF/AMP epithelium carried a large cytoplasmic vacuole, with apically displaced nucleus that was often crescent-shaped due to the vacuole (Fig. 2C). Nuclear FLTP-H2B-Venus expression indicated that all vacuolated cells were MCCs (Fig. 2D). We measured the distance between the base of the nucleus and the stroma (Fig. 2E) to quantify the severity of nuclear displacement and found that the apical displacement of MCC nuclei in 18mo mice was significantly larger than in 4mo mice (Fig. 2F). The nucleus of secretory cells (SCs) was also apically displaced and protruded from the epithelial monolayer in 18mo oviducts, although their nuclear shape remained round or oval (Fig. 2C, D). Similar to MCCs, the apical displacement of SC nuclei at 18mo was significantly larger than at 4mo (Fig. 2G). It is reported that SC protrusion from the epithelial monolayer is observed during the female hormonal cycle (17). In agreement with this, we found that protrusion of SC nuclei, marked by PAX8, was prevalent during diestrus stage (Fig. 2H, J), as compared to estrus stage (Fig. 2I, J) in young females, suggesting that SC protrusion was not exclusive to aging.

**Figure 2:**
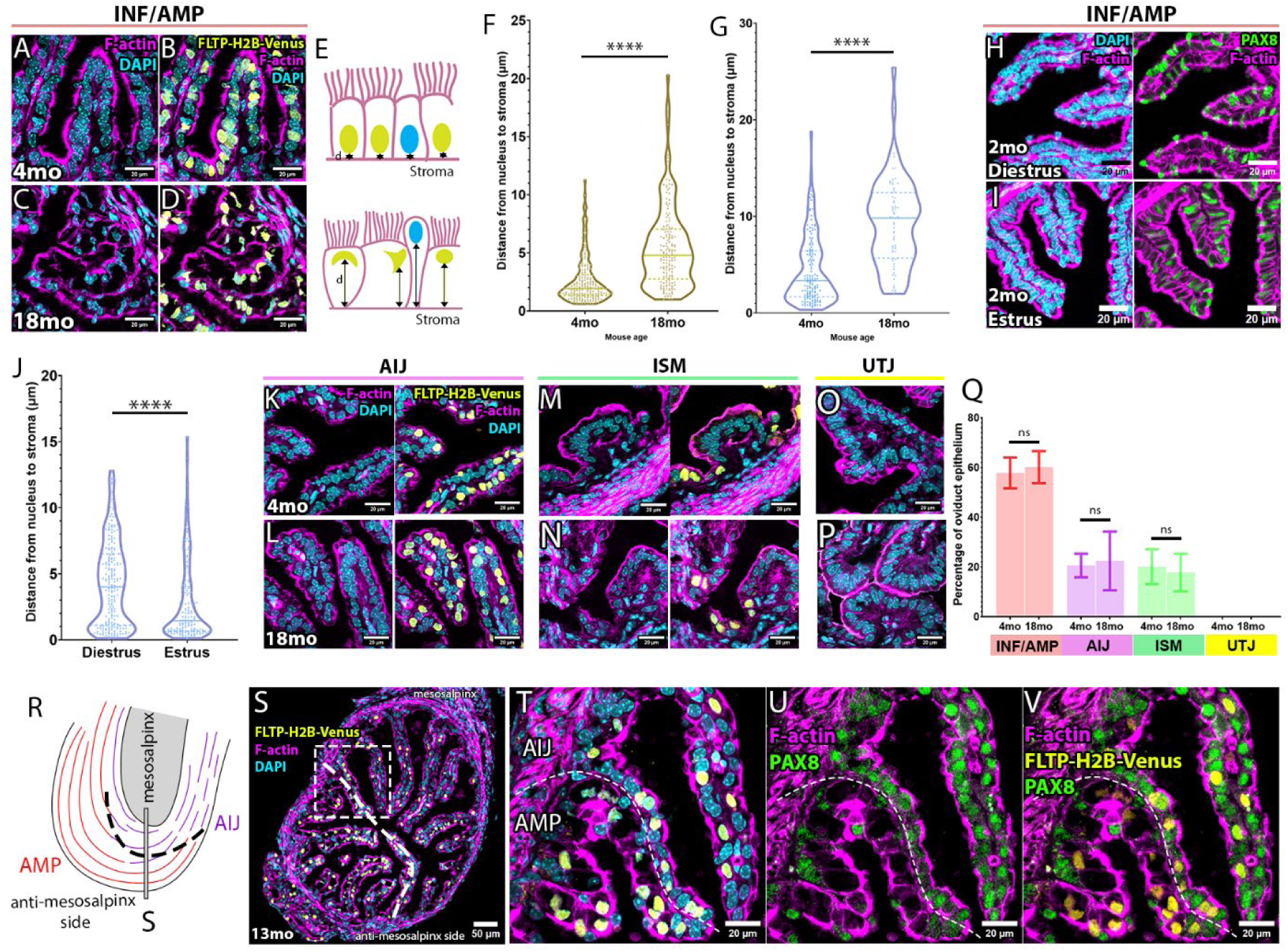
Aging-associated MCC vacuolation and accompanied apical displacement of the nucleus with nuclear anomaly are restricted in the INF/AMP epithelium. (A-D) Morphology of epithelial cells lining the INF/AMP mucosal folds in oviducts dissected from 4mo (A, B) and 18mo (C, D) mice. DAPI staining visualized the nuclear shape of epithelial cells, and FLTP-H2B-Venus marked MCC nuclei. In the 18mo INF/AMP epithelium, MCC nuclei were apically displaced (D), relative to 4mo INF/AMP (B). (E) Illustration showing measurement of nuclear apical displacement in 4mo and 18mo INF/AMP epithelium. (F) Measurements of the distance between MCC nuclei to stroma in the INF/AMP region of 4mo (mean ± SD, 2.606 ± 1.958 µm; N=3) and 18mo (5.482 ± 3.541 µm; N=3) oviducts. (G) Measurements of the distance between SC nuclei to stroma in the INF/AMP region of 4mo (4.415 ± 3.45 µm; N=3) and 18mo (9.324 ± 4.559 µm; N=3) oviducts. Each point represents a measurement/cell. The continuous line represents the median, while the dotted lines show the quartiles. **** = p < 0.0001. (H, I) Morphology of the INF/AMP epithelium of young 2mo mice in estrus stage. PAX8+ve SCs were aligned with other epithelial cells in the epithelium during estrus (I), while they apically protruded from the epithelium during diestrus (H). (J) Measurements of the distance of SC nuclei to stroma in the INF/AMP region of mice in estrus stage (2.791 ± 3.007 µm; N=3) and in diestrus stage (4.28 ± 3.219 µm; N=3) oviducts. (K-P) Morphology of epithelial cells lining the AIJ (K, L), ISM (M, N) and UTJ (O, P) mucosal folds in oviducts dissected from 4mo (K, M, O) and 18mo (L, N, P) mice. (Q) Percentage of MCCs in the INF/AMP, AIJ, ISM and UTJ of 4mo and 18mo oviducts, ns = p > 0.05. Error bars indicate standard deviation. (R) Illustration of the bevelled, sharp boundary of mucosal fold morphology, cell subtype, and change in proportion of MCCs between turns 2 and 3. (S) Transverse section of the sharp boundary between turns 2 and 3, as illustrated in (R). Sudden drop in MCC proportion on the mesosalpinx side, characteristic of the AMP-AIJ boundary. (T-V) Apical displacement of nuclei restricted to the AMP, but not evident in the AIJ (T). Change in PAX8 expression in AMP and AIJ epithelium (U), with MCCs being PAX8-ve in the AMP but PAX8+ve in the AIJ (V), confirming distinct cell subtypes at this boundary.

Interestingly, the MCC vacuolation and apically displaced crescent-shaped nuclei were observed in the INF/AMP epithelium (Fig. 2A-D) but not in the other regions of the oviduct. The nuclear shape in the AIJ, ISM and UTJ epithelium was generally round or oval, with no sign of cytoplasmic vacuolation (Fig. 2K-P). There was no apparent difference in the distribution pattern (Fig. 2A-D, K-P) and proportion (Fig. 2Q) of MCCs between 4mo and 18mo oviducts. We concluded that MCC vacuolation and apical displacement of the nucleus were aging phenotypes unique to the INF/AMP epithelium.

We previously established that there is a sharp boundary between each distinct region of the oviduct, visualized by changes in transcription factor expression, MCC proportion/distribution and mucosal fold morphology (1). At the bevelled AMP-AIJ boundary, located between oviduct coiling/turning points 2 and 3, the AMP epithelium occupied the anti-mesosalpinx side recognized by a higher proportion of MCCs, while the AIJ epithelium began on the mesosalpinx side (Fig. 2R, S). We found that only AMP MCCs, indicated by lack of PAX8 expression, had a large cytoplasmic vacuole with an apically displaced, crescent-shaped nucleus, while the AIJ MCCs did not show any apparent morphological abnormality, despite identical proximity to the ovary (Fig. 2T-V). This suggests that the aging-associated MCC vacuolation in the INF/AMP epithelium is linked with its unique cell property rather than physical proximity to the ovary.

### Progressive increase in size, frequency of MCC vacuolation and apical nuclear displacement in the INF/AMP epithelium after 6 months

To determine the timeline of this age-related phenotype in the distal INF/AMP epithelium, we examined 7mo and 13mo oviducts. Both 7mo and 13mo INF/AMP epithelium had varying proportions of apically displaced MCC nuclei, with most SC nuclei protruding from the epithelial monolayer (Fig. 3A-D). The position of the nucleus in INF/AMP MCCs was progressively apically displaced from 4mo to 18mo (Fig. 3C). Apical displacement of SC nuclei at 13mo was significantly larger than 4mo and 7mo, but not significantly different from 18mo (Fig. 3D), due to SC body protrusion from the epithelial monolayer at 13mo and 18mo (Fig. A, B). We found individual female variation in 7mo oviducts: 1) some individuals exhibited exaggerated apical displacement of MCC nuclei and large, empty cytoplasmic spaces in many cells (Fig. 3E; N=2/3), similar to that noted in 13mo or 18mo mice, and 2) an individual presented with low-to-moderate apical displacement with small cytoplasmic spaces (Fig. 3F; N=1/3).

**Figure 3:**
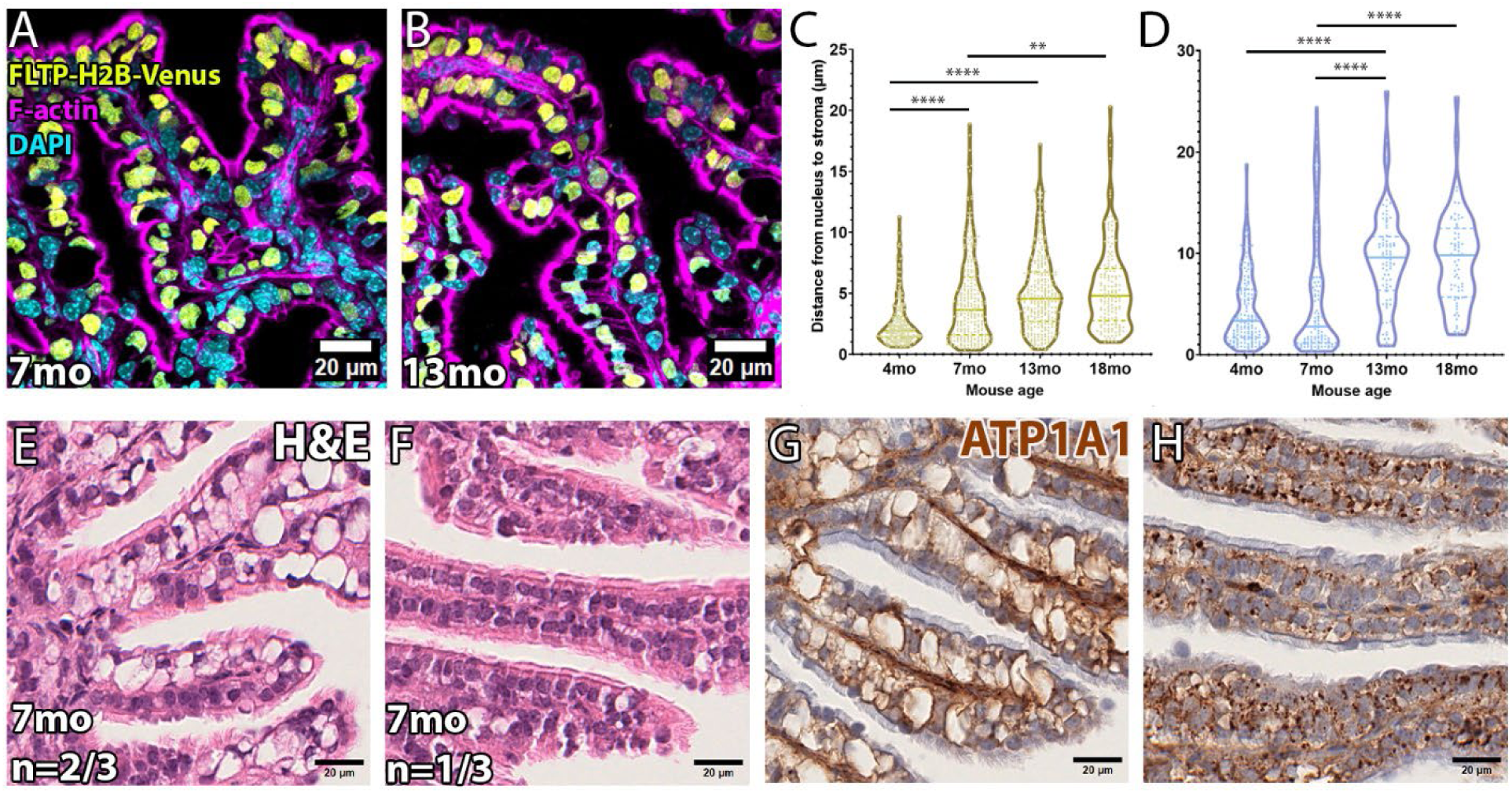
Progressive apical displacement of the nucleus and large vacuolation with aqueous fluid in INF/AMP MCCs. (A, B) Morphology of epithelial cells lining the INF/AMP mucosal folds in oviducts dissected from 7mo (A) and 13mo (B) mice. SC nuclei and deformed MCC crescent-shaped nuclei were apically displaced. (C) Measurements of the distance of MCC nuclei to stroma in the INF/AMP region of 4mo, 7mo (4.476±3.543 µm; N=3), 13mo (5.014±3.085 µm; N=3), and 18mo oviducts. (D) Measurements of the distance of SC nuclei to stroma in the INF/AMP region of 4mo, 7mo (5.628±6.045 µm; N=3), 13mo (9.285±4.790 µm; N=3), and 18mo oviducts. Each point represents a measurement/cell. The continuous line represents the median, while the dotted lines show the quartiles. **** = p < 0.0001, ** = p < 0.001. (E, F) H&E staining of the 7mo INF/AMP region. Two phenotypes were noted: 1) exaggerated/large, empty cytoplasmic vacuoles and apically displaced nuclei (E) and 2) small/indiscernible empty spaces/vacuoles and nuclei located in the centre of the cells (F). (G, H) ATP1A1 expression in 7mo INF/AMP region. Large cytoplasmic vacuoles were lined with ATP1A1 (G), while MCCs with small/indiscernible vacuoles contained ATP1A1+ve punctae (H).

A vacuole was located at the basal side of MCCs, clearly distinct from secretory vesicles typically located at the apical side. The vacuole appeared to cause apical displacement and crescent-shaped deformation of the nucleus. Haematoxylin and eosin (H&E) staining showed empty cytoplasmic spaces, ruling out the possibility of diluted cytoplasm, protein inclusions, or nucleic acid accumulation (Fig. 3E, F). In agreement with this, a classic TEM study of the AMP epithelium in 7-to 24-month-old mice show MCCs with large vacuoles containing heterogeneously dense materials, with no obvious changes in other organelles including cilia (17). We found that the large cytoplasmic vacuole was lined with sodium/potassium-transporting ATPase subunit alpha-1 (ATP1A1; Fig. 3G) and did not show lipid or glycogen accumulation (Suppl. Fig. 1A-C), suggesting aqueous/fluid accumulation in INF/AMP MCCs. Cytoplasmic, punctate ATP1A1 localization surrounded the small vacuoles in MCCs with low-to-moderate apical displacement (Fig. 3H). Cytoplasmic ATPA1 was not observed in epithelium of the proximal regions (Suppl. Fig. 1D-F). Occasional punctate ATP1A1 was noted in the INF/AMP of 3mo mice (Suppl. Fig. 1G).

### INF/AMP MCC vacuolation was independent of ovulation and estrous cycling

The INF/AMP region is located directly adjacent to the ovary, a reproductive organ that undergoes ovulation every 4-5 days in mice (18). We considered two possibilities: 1) the distal oviduct is exposed to events associated with ovulation, including exposure to oxidative stress, inflammation-like response, and production of reactive oxygen species (ROS) that induce repetitive stress in the adjacent INF/AMP epithelium (19), and 2) the other possibility is estrous cycle-associated events like epithelial cell turnover. It is known that the INF/AMP epithelium shows fluctuation in the proportion of SCs and MCCs in response to hormonal regulation (20). Decline in estrous cyclicity occurs by ∼9-months in inbred mice (21), characterized by a progressive lengthening of estrous cycles followed by cyclic cessation and anestrus by 11 to 16-months of age. Thus, age-associated lack of the hormonal cycling may contribute to MCC vacuolation due to reduced cell turnover.

To address these possibilities, we performed bilateral ovariectomy (OVX) in 2mo mice and evaluated the short and long-term impact to oviduct epithelial cells. For short-term analysis, we isolated the reproductive tracts 1-month post OVX. In these oviducts, discernible cytoplasmic vacuoles were not observed (Fig. 4A), suggesting that MCC vacuolation was not a response to the lack of the estrous cycling. It is of worth to note that the INF/AMP SC nuclei were displaced apically due to SC protrusion from the epithelial monolayer (9.124±3.146 µm; Fig. 4A), similar to the diestrus stage in non-OVX females.

**Figure 4:**
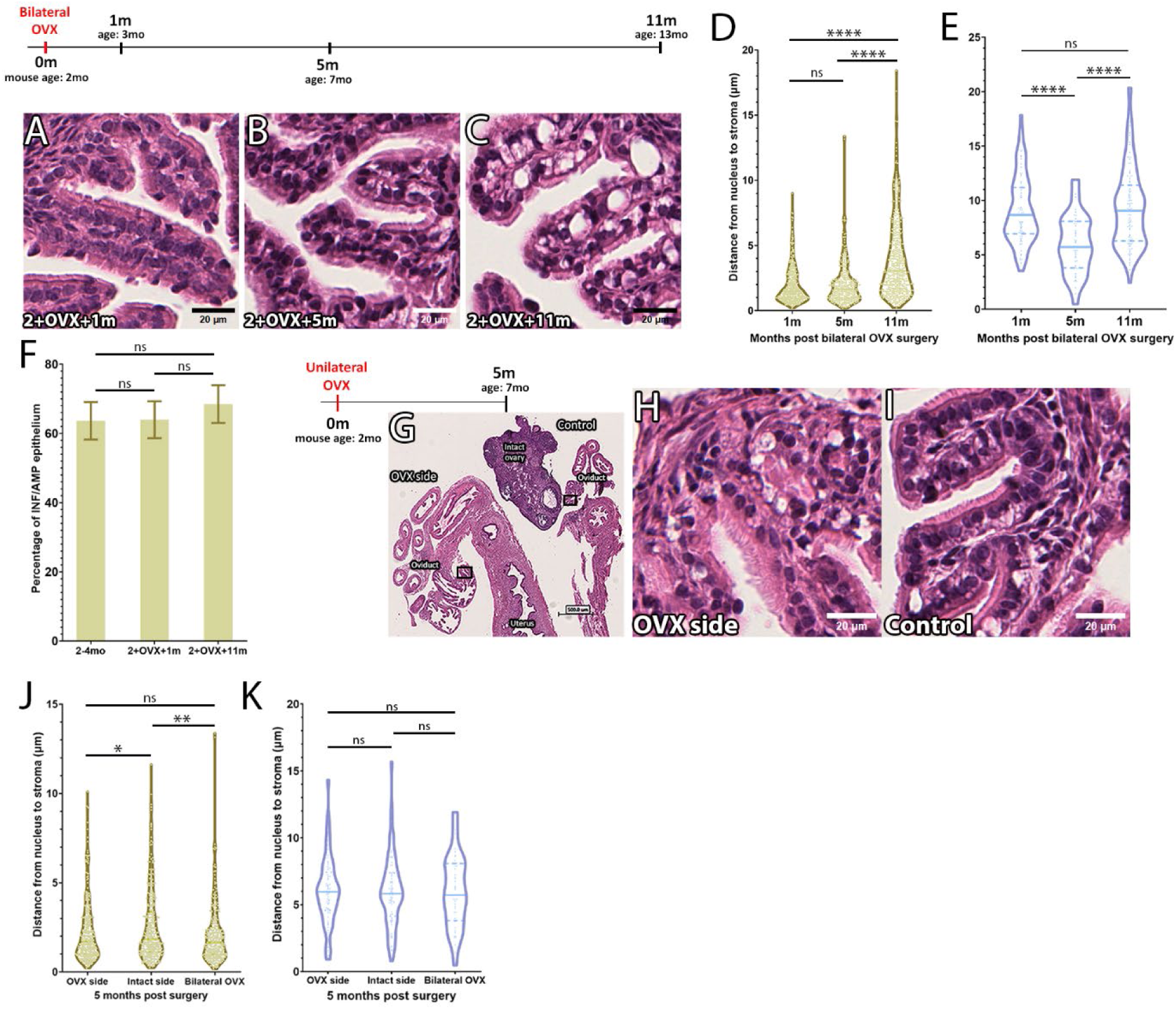
Timeline of the INF/AMP MCC vacuolation is unaffected regardless of loss of ovulation and estrous cycling. (A-C) H&E staining of the INF/AMP region in oviducts dissected from mice 1 month (A), 5 months (B), and 11 months (C) following bilateral OVX. MCC vacuolation are observed in the INF/AMP epithelium of mice 5– and 11-months post bilateral OVX. (D) Measurements of the distance of MCC nuclei to stroma in the INF/AMP epithelium of mice 1m (1.937±1.402 µm; N=3), 5m (2.077±1.656 µm; N=3), and 11m (3.977±2.977 µm; N=4) post bilateral OVX. (E) Measurements of the distance of SC nuclei to stroma in the INF/AMP epithelium of mice 1m (9.124±3.146 µm), 5m (5.994±2.682 µm), and 11m (9.334±3.694 µm) post bilateral OVX. Changes in apical displacement of SC nucleus indicated corresponded with changes in the height of the epithelium. (F) Percentage of MCCs in the INF/AMP epithelium of oviducts from non-OVX 2-4mo mice, and OVX mice dissected at 1m and 11m post surgery, ns = p > 0.05. Error bars indicate standard deviation. (G-I) H&E staining of the INF/AMP region in oviducts dissected from mice 5 months following unilateral OVX. MCC vacuolation was noted in both the ovariectomized/OVX side (H) and the side with the ovary intact/intact side (I). (J) Measurements of the distance from MCC nuclei to stroma in the INF/AMP region of mice 5m following unilateral OVX, comparing distances on the OVX side (2.220±1.705 µm; N=3) and intact side (2.550±1.982 µm; N=3) to the INF/AMP region of mice 5m following bilateral OVX. The intact side had the largest apical nuclear displacement. (K) Measurements of the distance of SC nuclei to stroma in the INF/AMP region of mice 5 months following unilateral (OVX side = 6.013±2.493 µm, intact side = 5.955±2.642 µm) and bilateral OVX. Each point represents a measurement/cell. The continuous line represents the median, while the dotted lines show the quartiles. **** = p < 0.0001, ** = p < 0.001, * = p< 0.01, ns = p > 0.05.

For the long-term analyses, we isolated reproductive tracts 5– and 11-months post OVX. At 5-months post OVX, when the age of these mice was 7mo, we observed MCC vacuolation in the INF/AMP epithelium (Fig. 4B), showing that the timeline of MCC vacuolation was unaffected by OVX. In agreement with this, vacuoles were not noted in the INF/AMP of mice 3-months post OVX, when the age of the mice was 5mo (Suppl. Fig. 2A). Thus, OVX did not facilitate or prevent MCC vacuolation. Interestingly, the apical displacement of the nucleus in MCCs at 5-months post OVX was not significantly different from 1-month post OVX (Fig. 4D), despite noticeable vacuole formation. Instead, the cellular height of the epithelium in mice 5-months post OVX (8.557±1.84 µm) was lower than 1-month post OVX (9.427±2.244 µm). The shape of MCCs was cuboidal with a round nucleus in OVX females while it is packed and columnar with an oval-shaped nucleus in non-OVX females. Taken together, bilateral OVX did not prevent or facilitate MCC vacuolation, but the height of the epithelial cells was decreased. This was reflected in a significant decrease in the apical displacement of INF/AMP SC nuclei in 5-months post OVX despite obvious protrusion from the epithelial monolayer (Fig. 4B, E). A large cytoplasmic vacuole was noted in individual INF/AMP MCCs at 11-months post OVX, with an apically displaced nucleus (Fig. 4C, D). INF/AMP SC nuclei at 11-months post OVX were also apically displaced relative to 5-months post OVX, due to progressive enlargement of cytoplasmic vacuoles resulting in increased epithelial cell height at 11-months post OVX. Finally, despite cell shape change and loss of estrous cycling/ovulation, there was no significant difference in the proportion of MCCs in the INF/AMP region isolated from mice that were 2-4mo, or 1– and 11-months post bilateral OVX (Fig. 4F), suggesting low epithelial cell turnover, consistent with previous studies (20, 22). No discernible cytoplasmic vacuoles were noted in the AIJ, ISM, and UTJ at 11-months post OVX (Suppl. Fig. 2B-D).

Since bilateral OVX completely terminates the ovarian hormonal cycle, we performed unilateral ovariectomy surgery in 2mo mice to delineate ovulation and hormonal cycle-associated homeostatic events in the epithelium. We isolated reproductive tracts 5-months post unilateral OVX (Fig. 4G) and found MCC vacuolation on both sides, with no apparent difference between OVX and control sides (Fig. H, I). Maintenance of estrous cycling without ovulation did not prevent MCC vacuolation. However, apical displacement of MCC nuclei on the OVX side was smaller than the control/intact side, likely due to cyclic cell shape changes to accommodate OCCs following ovulation. Additionally, there was no significant difference in MCC nucleus apical displacement between the OVX side and mice 5-months post bilateral OVX (Fig. 4J) or in displacement of SC nuclei (Fig. 4K). Thus, we conclude that the MCC vacuolation is not caused by repetitive ovulation or estrous cycle-associated events.

### Enrichment of genesets associated with ER, metabolic, mitochondrial stress and/or ATP metabolism in aged INF/AMP cells, suggesting a metabolically stressed microenvironment

To investigate the changes in the aged INF/AMP microenvironment accompanying MCC vacuolation, we performed single-cell RNASeq of 16,948 cells pooled from the INF/AMP region of three 18mo mice (Aged_INFAMP) and compared this to 10,281 cells isolated from the INF/AMP of a single estrus stage 3mo mouse (Estrus_INFAMP). Cells from the INF/AMP region of the mouse oviduct clustered into 15 clusters that were then grouped into 10 cell types/populations based on a combination of their top 5 highly expressed genes and known cell subtype markers (Fig. 5A, Suppl. Fig. 3, 4A-M). These cell populations include: 1 –secretory cells/SCs (cluster 1, 2, 11 expressing Epcam, Wt1, Pax8, Ovgp1, Sox17; Suppl. Fig. 3, 4A-D), 2 –stromal cells (cluster 3, 7, 10 expressing Dcn, Igfbp6, Pdgfra; Suppl. Fig. 3, 4F), 3 –multi-ciliated cells/MCCs (cluster 4 expressing Epcam, Wt1, Foxj1; Suppl. Fig. 4A, B, E), 4 –smooth muscle cells (cluster 5, 14 expressing Acta2, Actg2, Dmd; Suppl. Fig. 3), 5 –endothelial cells (cluster 6 expressing Pecam1, Sox17; Suppl. Fig. 4D, G), 6 –antigen presenting cells/APCs (cluster 8 expressing Cd52, H2-Aa; Suppl. Fig. 3, 4H), 7 –vascular smooth muscle cells/VSMCs (cluster 9 expressing Acta2, Rgs5; Suppl. Fig. 3), 8 –T-cells (cluster 12 expressing Cd52, Cd3e; Suppl. Fig. 4H, I), 9 –mesothelial cells (cluster 13 expressing Wt1, Msln, Dcn, Igfbp6; Suppl. Fig. 3, 4B, H), and 10 –red blood cells/RBCs (cluster 15 expressing Hbb-bs, Suppl. Fig. 3). Surprisingly, despite the apparent morphological differences, there was no difference in cell subtypes and populations. All populations in Estrus_INFAMP and Aged_INFAMP overlapped with each other (Fig. 5B). In all clusters, cells from Aged_INFAMP had more upregulated genes than downregulated genes, relative to Estrus_INFAMP (Fig. 5C).

**Figure 5:**
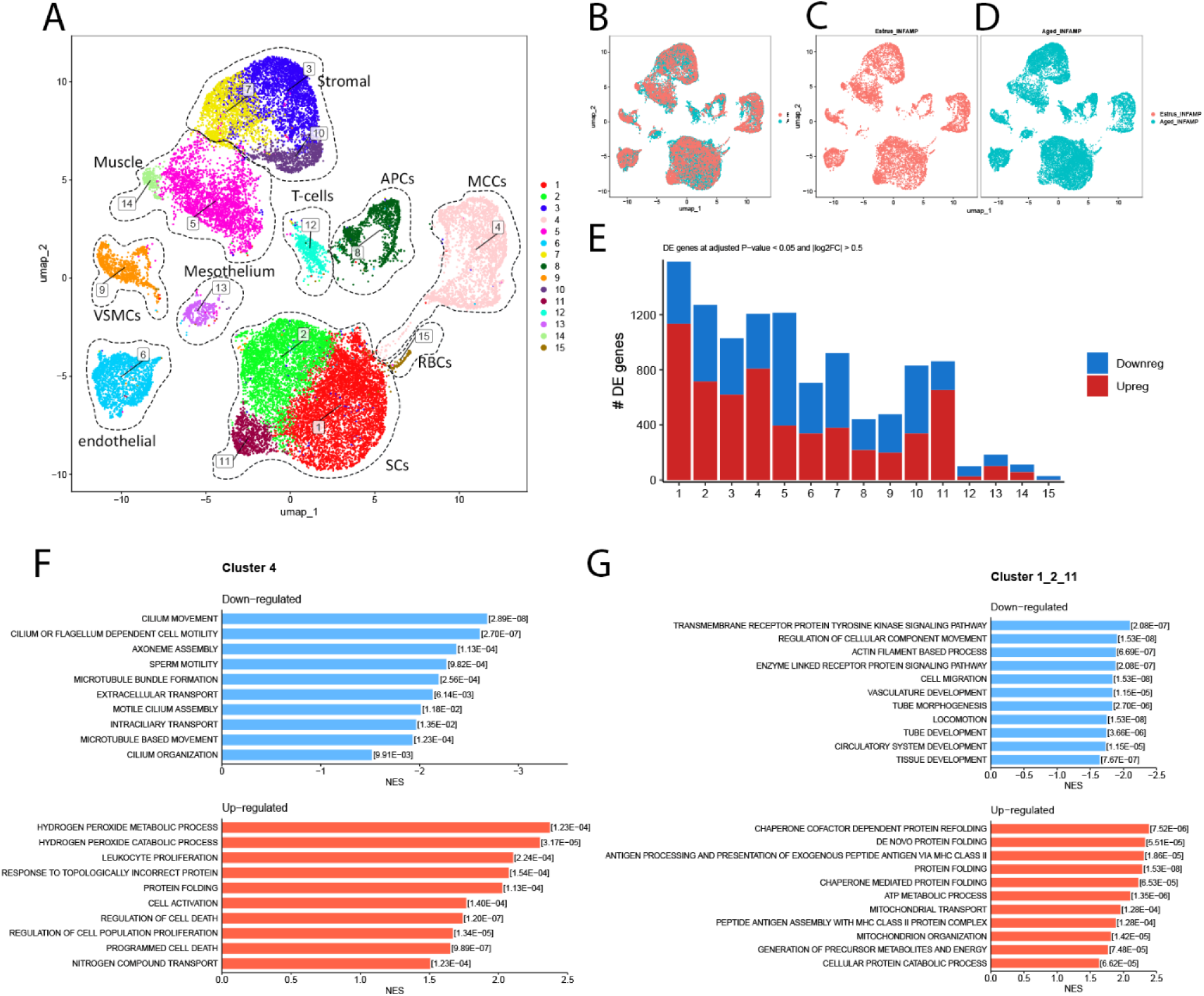
ROS metabolism, ER stress, and cell death regulation gene-sets were upregulated in 18mo INF/AMP MCCs, while ER stress, ATP metabolism, mitochondrial transport gene sets were upregulated in 18mo INF/AMP SCs. (A) Integrated UMAP plot colour-coded by cluster membership, cell populations are labelled and enclosed within dotted lines, identified based on top 5 genes and known markers (A). (B-D) UMAP clustering of cells isolated from 3mo estrus stage mouse (C), 18mo mice (D), and overlap between the two (B) showing no significant alteration in the isolated populations. (E) Number of differentially expressed genes in each cluster, adjusted for p-value < 0.05 and |log2FC| > 0.5, between INF/AMP cell populations isolated from 3mo estrus stage mouse and 18mo mice. (F) Gene sets downregulated (top, blue) and upregulated (bottom, orange) in INF/AMP MCCs of 18mo mice, as compared to a 3mo estrus stage mouse. (F) (G) Gene sets downregulated (top, blue) and upregulated (bottom, orange) in INF/AMP SCs of 18mo mice, as compared to a 3mo estrus stage mouse.

Using Gene Set Enrichment Analysis (GSEA), we found that aged 18mo INF/AMP MCCs showed downregulation of genes associated with cilia movement and organization, and upregulation of genes associated with hydrogen peroxide metabolism, unfolded protein response (UPR), and negative regulation of cell death (Fig. 5D). This reflects an adaptive/compensatory response to cellular stress, in particular, endoplasmic reticulum (ER) stress in MCCs. Further, aged 18mo INF/AMP SCs showed downregulation of genes associated with transmembrane receptor protein tyrosine kinase signaling, actin filament-based process, and cell migration/movement. Similar to the MCCs, they showed upregulation of genes associated with stress responses such as UPR, ATP metabolic process, and mitochondrial organization/transport (Fig. 5G; Suppl. Fig. 5A-C).

Stromal cells were divided into three clusters and showed downregulation of genes associated with ECM production in cluster 3 and 7 (Suppl. Fig. 5D, E), ribosome biogenesis and RNA processing in cluster 10 (Suppl. Fig. 5F). Cluster 3 stromal cells showed upregulation of gene sets associated with various biosynthetic and metabolic processes, negative regulation of cell death, and immune system activity regulation, reflecting adaptive mechanisms in response to cellular stress and/or a gradual decline in metabolic efficiency (Suppl. Fig. 5D). On the other hand, cluster 7 stromal cells did not show significant upregulation of gene sets, as compared to 3mo cells (Suppl. Fig. 5E). Cluster 10 stromal cells showed upregulation of genes associated with acute inflammatory response, cell adhesion, and cytokine production (Suppl. Fig. 5F), reflecting a response to inflammation/resolution of inflammation, facilitation of tissue repair, and/or cellular senescence.

Cluster 5 smooth muscle cells showed downregulation of genes associated with muscle organisation, blood circulation, RNA metabolic processes, and ribosome biogenesis, and upregulation in tissue growth, cell adhesion, and negative regulation of cell death (Suppl. Fig. 5G). There was no enrichment of genesets in cluster 14 smooth muscle cells (Suppl. Fig. 5I). Taken together, most populations were typically responding to cellular metabolic stresses, with a shift/decrease in biogenesis activities, and facilitation of tissue repair to maintain structural integrity. VSMCs showed downregulation of genes associated with tissue growth, while upregulation in genesets associated with vascular tone alteration, regulation of IGFR signaling, removal of ROS/RNS species, ATP metabolism, and monocarboxylic, carbohydrate metabolism (Suppl. Fig. 5H).

Among immune cells, T-cells and Antigen Presenting Cells (APCs: B-cells, macrophages, dendritic cells) showed upregulation of genesets associated with UPR (Suppl. Fig. 5J, K), suggestive of response to cellular stress. Endothelial cells showed downregulation in tissue and blood vessel growth, and upregulation in antigen presentation, superoxide radical/hydrogen peroxide removal, and various metabolic processes (Suppl. Fig. 5L). Finally, the mesothelial cells that line the INF/AMP serosa showed downregulation in nerve function and ECM organization, while upregulation in UPR, mitochondrial transport/organization, ATP metabolism, and regulation of cell death (Suppl. Fig. 5M), suggesting ER stress, energy demand compensation, and response to cellular stress.

Taken together, the single-cell transcriptomes of all isolated cells from 18mo INF/AMP indicated a general reduction of biogenesis and growth-related genes, and upregulated stress response genes and ATP energy demand compensation, without a clear indication of inflammation (Suppl. Fig. 5O, P).

### Hypoxia or hydroxyurea treatment induced MCC vacuolation *in vitro* organotypic slice cultures

It has been shown that cellular vacuolation in *in vitro* cultures occurs in response to ER stress, ATP depletion/shortage, cellular senescence, metabolic imbalance, bacterial toxins/viral components, exposure to drugs/toxic chemicals/pollutants, and hypoxia (23–25). To determine the potential cause of MCC vacuolation, we developed an *in vitro* organotypic slice culture system (150µm thick slices, see methods). Since many of the upregulated genesets in 18mo MCCs and SCs were associated with ER stress (Fig. 5E, F), we treated INF/AMP slice cultures with 5µM tunicamycin (TUN) for 24 hours to induce ER stress. However, TUN treatments did not result in any noticeable cellular changes (Suppl. Fig. 5A, B), suggesting that MCC vacuolation was not a consequence of ER stress.

ATP depletion using a combination of Antimycin A and 2-deoxyglucose (2-DG) treatment results in internalization of Na, K-ATPase in MDCK cells (26). The combined treatments of these two inhibitors visibly affected INF/AMP MCCs after 24hrs. Many MCCs protruded from the epithelial monolayer, along with condensed nuclei and occasional cleaved caspase 3 expression (Fig. 6A-E), suggesting apoptotic MCCs. While Antimycin A treatment alone also resulted in similar MCC protrusion in INF/AMP MCCs (Fig. 6G, J) and termination of cilia beating (data not shown), 2-DG alone did not (Suppl. Fig. 5C-E), suggesting that INF/AMP MCCs were highly dependent on cellular respiration. Antimycin A-induced MCC protrusion was unique to INF/AMP slices but not observed in AIJ and ISM slices (Fig. 6G-L), suggesting region-specific cellular respiration dependency. This is consistent with a recent study that showed distinct responses to mitochondrial dysfunction in the lung, dependent on cellular context (27).

**Figure 6:**
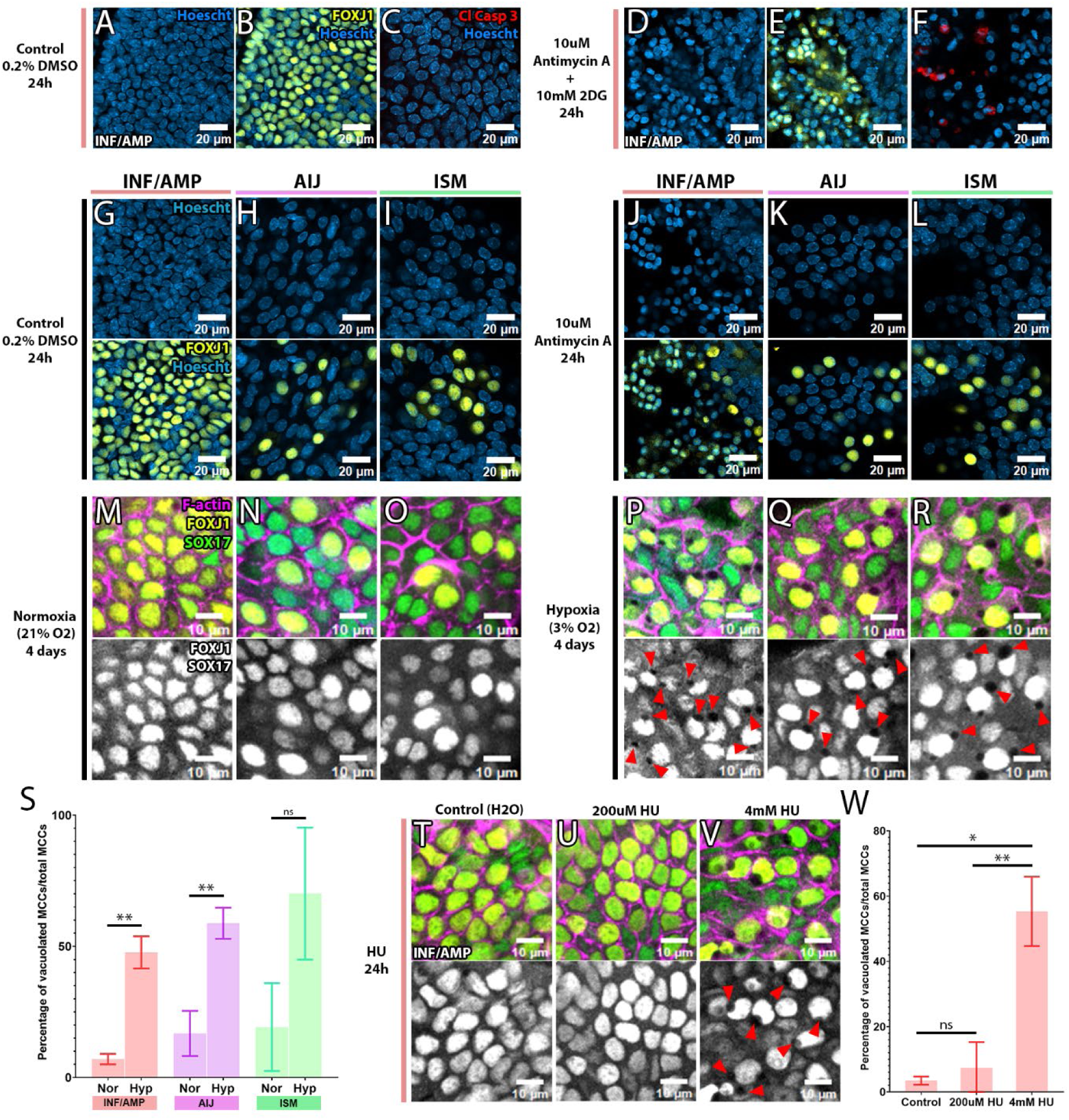
Hypoxia or HU treatment induced MCC vacuolation. (A-F) Inhibition of cellular respiration with 10μM Antimycin A and glycolysis with 10mM 2DG in organotypic slice cultures of the INF/AMP region. (A-C) Control cultured slice from INF/AMP region with round nuclei (A), of which most cells were FOXJ1+ve MCCs (B) and Cleaved Caspase 3-ve, suggesting no apoptotic cells (C). (D-F) Treatment with Antimycin A and 2-DG for 24 hours resulted in MCCs with high intensity nuclear staining (D, E), of which some cells were positive for Cleaved Caspase 3, suggesting apoptotic cells (F). (G-L) Inhibition of cellular respiration in organotypic slice cultures of the INF/AMP (G, J), AIJ (H, K), and ISM (I, L). 24-hour-long Antimycin A treatment resulted in MCC protrusion from the epithelial monolayer and high intensity of nuclear staining (J) but did not affect AIJ and ISM MCCs (K, L). (M-X) Organotypic slice culture for 4 days in normoxia (M-R) or hypoxia (S-X). When cultured in hypoxia, FOXJ1+ve MCCs of the INF/AMP (P), AIJ (Q) and ISM (R) present with empty vacuoles and deformed nucleus, while very few vacuoles formed in normoxia (M-R). FOXJ1 and SOX17, marking oviduct MCCs and SCs, were changed to white for easier visualization of vacuoles (P-R, V-X). Red arrows mark the vacuoles. (Y) Percentage of vacuolated MCCs in the INF/AMP, AIJ, and ISM in normoxia (7.005±1.995%, 16.971±8.603%, 19.205±16.739% respectively) versus hypoxia (47.728±6.129%, 58.82±5.938%, 60.283±28.439% respectively). Increased instances of vacuolation were noted in hypoxia, as compared to normoxia. Error bars indicate standard deviation. ** = p < 0.001. (Z-e) Hydroxyurea (HU) treatment for 24 hours in INF/AMP organotypic slice cultures. 200µM/low dose HU treatment did not result in discernible cellular changes (a, d), while 4mM/high dose HU treatment resulted in cytoplasmic vacuolation and nucleus deformation (b, e). FOXJ1 and SOX17, marking oviduct MCCs and SCs, were changed to white for easier visualization of vacuoles (c-e). (f) Percentage of vacuolated MCCs in the INF/AMP when treated with water (2.786±0.573%), low dose HU (7.397±7.865%), and high dose HU (55.32±10.622%) for 24 hours. Increased instances of vacuolation were noted in high dose HU treatment, as compared to low dose or water control. Error bars indicate standard deviation. ** = p < 0.001, ** = p < 0.001, ns = p > 0.05.

No significant enrichment in gene sets associated with cellular senescence was noted in the single-cell transcriptomic analysis (Suppl. Fig. 5F). Although we found the absence of Lamin-B1, often associated with cellular senescence, in the 18mo oviduct (Suppl. Fig. 5G-J), we discovered that Lamin-B1 showed a cyclic expression pattern during the estrous cycle in young females, with high intensity nuclear membrane localization in the estrus stage (Suppl. Fig. 5K) and low intensity in diestrus stage (Suppl. Fig. 5L). We did not find significant changes in the expression of cellular senescence-associated pseudo-DNA damage response genes including cell cycle arrest-associated p21 and p16, or DNA damage marker phospho-H2Ax (28) in 18mo and 2mo mice (Suppl. Fig. 5M-R). Unexpectedly, phospho-H2Ax nuclear puncta were predominantly observed in INF/AMP MCCs throughout adulthood and in aging (Suppl. Fig. 5Q, R).

Induction of metabolic stress/nutrient deprivation with 2µM 6-AN (pentose phosphate pathway inhibitor; Suppl. Fig. 5S), 0.8µM ST045849 (hexosamine biosynthesis pathway inhibitor; (Suppl. Fig. 5T), 1µM YZ9 (PFKFB3/glycolysis inhibitor; Suppl. Fig. 5U), 1µM Shikonin (PKM2/glycolysis inhibitor; Suppl. Fig. 5V), and 10µg/ml CHX (cycloheximide, protein synthesis inhibitor; Suppl. Fig. 5W) for 24 hours in *in vitro* organotypic slice cultures did not result in significant, discernible cellular changes.

Although there was no significant enrichment of gene sets associated with hypoxia in 18mo INF/AMP (Suppl. Fig. 5X), *in vitro* INF/AMP tissue slices cultured for 4 days in 3% O_2_ (chronic, mild hypoxia) reproducibly induced a cytoplasmic vacuole and deformed nucleus in MCCs (Fig. 6P, S), but similar 4-day-long culture in 21% O_2_ (normoxia) did not (Fig. 6M). Chronic hypoxia results in reduced ATP demand and rate of mitochondrial oxygen consumption due to ROS elevation (29). Thus, the cytoplasmic vacuolation induced by hypoxia could be a MCC response to ATP shortage and not ATP depletion. Interestingly, hypoxia-induced MCC vacuolation was not restricted to the INF/AMP epithelium but also observed in AIJ and ISM epithelia (Fig. 6M-O, S). No vacuolated SCs were noted in slice cultures of any of the regions (Fig. 6M-R, Suppl. Fig. 5Y, Z). *In vitro* experiments suggest that MCCs are more sensitive to hypoxia than SCs, while the cause for unique MCC vacuolation in the INF/AMP epithelium *in vivo* remains in question.

HU treatment of *in vitro* organotypic slice cultures with low dose HU, known to induce ROS in cycling cells (30), did not result in discernible cellular changes (Fig. 6T, U, W). Interestingly, 24-hour-long treatment with high dose HU, known to cause dNTP depletion and replication stress in cycling cells (31), induced a large cytoplasmic vacuole with a deformed nucleus in MCCs (Fig. 6V, W).

### MCC vacuolation limited to the INF/AMP epithelium *in vivo* correlated to aging-associated changes in blood flow via the ovarian artery

Although our *in vitro* experiments indicated that all oviduct MCCs were sensitive to hypoxic stress and form cytoplasmic vacuoles, age-associated MCC vacuolation was uniquely restricted to the INF/AMP MCCs *in vivo*. This suggests that, in addition to the distinct cellular property of the INF/AMP epithelium (1, 13, 20), the local environment within the INF/AMP could be different from the rest of the oviduct. Since metabolic stress-associated gene sets were upregulated, we investigated blood circulation that supplies the reproductive tract. Three arteries feed the female reproductive organs: ovarian, uterine and vaginal arteries. The ovarian artery is derived from the abdominal aorta, while uterine and vaginal arteries are derived from the internal iliac artery (32, 33). These patterns are evolutionarily conserved between mice and humans despite anatomical differences in the female reproductive tract.

To track circulation, we performed cardiac perfusion of CellMask, a fixable cell membrane staining fluorescent dye. This allowed us to visualize the blood vessels around the female reproductive tract, particularly the ovarian and uterine arteries (Fig. 7A, B, F, G). We noticed that a branch of the ovarian artery fed into the ovary and parts of the oviduct (Fig. 7A, B, C-E, H-J), while the uterine artery fed into the proximal oviduct and the uterus (Fig, 7A, B, F, G). Interestingly, this branch of the ovarian artery ran along the mesosalpinx, branching into two vessels, with one directly feeding into the INF/AMP and the other supplying parts of the proximal oviduct (Fig. 7D, E, I, J). Since the INF/AMP was supplied by only this branch of the ovarian artery, we speculated that circulation to the ovary was linked to that of the INF/AMP. Further, using the *Flk1-gfp* mouse line, which labels endothelial cells at embryonic stages (34), we found an intricate network of vessels throughout the upper female reproductive tract, likely marking the microvasculature (Fig. 7K-M). FLK1-GFP+ve vessel/vascular density was sparse in the AMP, as compared to the ISM (Fig. 7N, O), suggesting distal-proximal differences in microvasculature. Interestingly, while CellMask entered the oviduct in young mice (Fig. 7C), indicating dilated vessels/active circulation, CellMask was not noted in the ovaries or oviducts of aged mice (Fig. 7P, Q), suggesting changes/decline in circulation. Circulation into the ovary declined in tandem with ovarian aging, correlating with reduction in ovary size and number of oocytes in aged mice (Fig. 7R-U). Ovarian aging and oocyte depletion progressively occurs in 3-12mo mice, with decrease in number of oocytes at 6mo, and pronounced reduction at 9mo and 12mo (35). This timeline of oocyte depletion and ovarian aging occurs concurrently with INF/AMP MCC vacuolation, likely due to changes in circulation via the ovarian artery.

**Figure 7:**
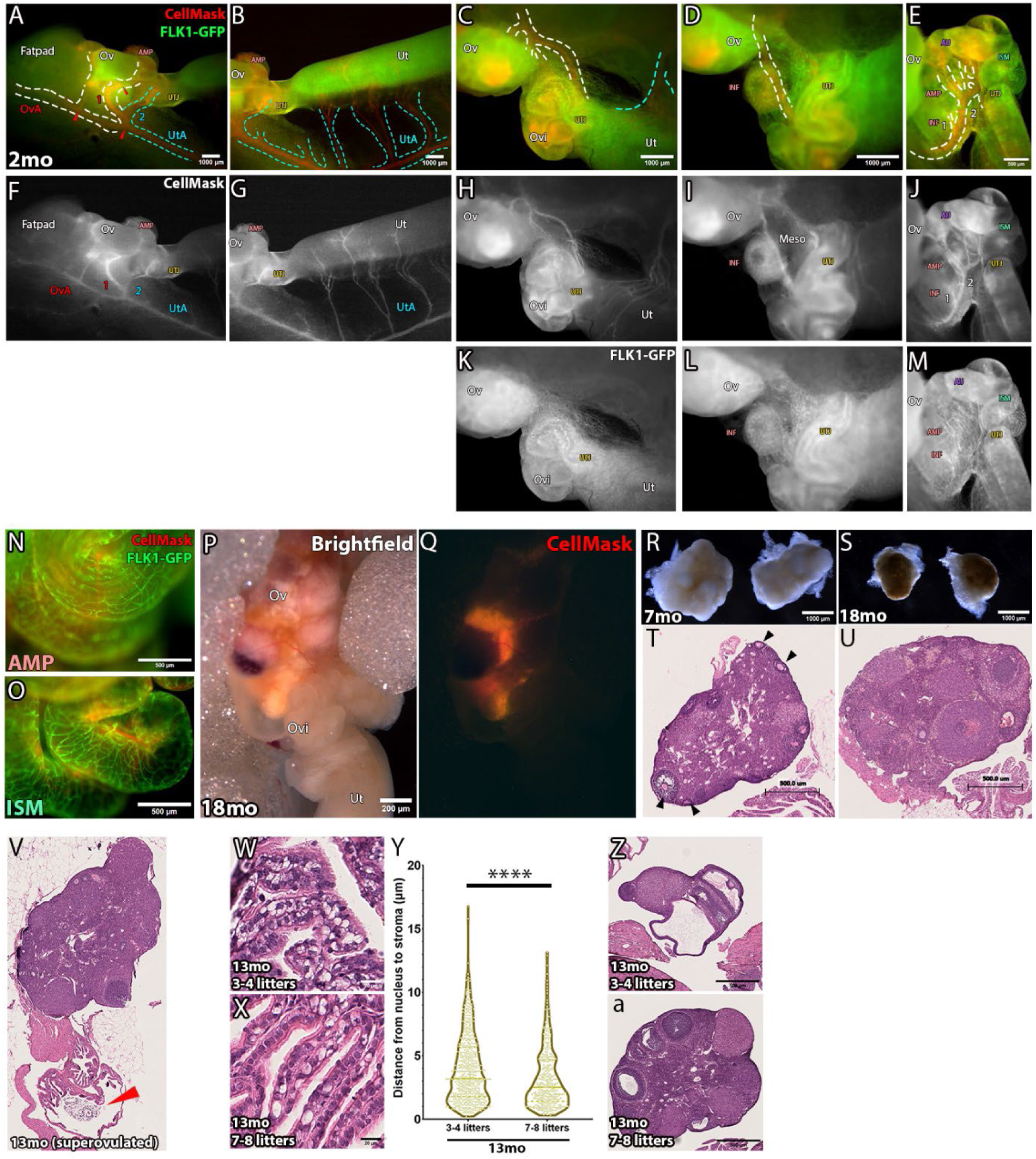
Reduced circulation via the ovarian artery in aged females. (A-J) A 2mo oviduct. Ovarian artery (OvA) and uterine artery (UtA) were labelled by cardiac perfusion of CellMask in a 2mo *Flk1-gfp* female. The ovarian artery ran along the connective tissue that connected the ovary (Ov), oviduct (Ovi) and uterus (Ut), with branches feeding into the ovary. From the anastomosis of the ovarian and uterine arteries, two branches diverged: 1) one fed into the ovary and branched again to supply the distal oviduct, and 2) the other fed into the proximal oviduct. The ovarian and uterine arteries were traced by white and cyan dotted lines, respectively. Branching is labelled by red arrows. (A, F). Following the anastomosis point, the uterine artery ran along the broad ligament on the mesometrial side, with numerous branches connecting into the uterus (B, G). The ovarian artery as seen from a different angle, with branches that connected into the ovary and the oviduct (C, H). The oviduct was partially stretched out to visualise a branch of the ovarian artery that ran along the mesosalpinx (D, I). This artery further branched out: 1) one branch directly fed into the INF/AMP, while 2) the other branch split into numerous vessels that supplied the proximal oviduct (E, J). (K-M) FLK1-GFP expressing vessels formed an intricate network throughout the upper reproductive tract, likely labeling arterioles and/or capillaries. (N, O) Microvasculature differences between distal and proximal oviduct. Network of FLK1-GFP expressing vessels was less dense in the AMP, relative to the ISM. (P, Q) Lack of CellMask labeling in in the upper female reproductive tract of aged 18mo mice. (R-U) Age-associated ovarian atrophy, decrease in ovary size, and oocyte numbers. Ovaries from 18mo mice were smaller and had a smooth surface (S) and no oocytes were noted in 18mo ovaries (U), as compared to 7mo ovaries (R, T). (V) MCC function was unaffected by vacuolation. OCCs were noted in the AMP region of 13mo superovulated mice. (W-Y) Reduced severity in apical displacement of nucleus in INF/AMP MCCs of 13mo mice with higher gravidity. Fewer vacuolated MCCs were noted in 13mo mice that have had 7-8 litters (X), relative to those that have had 3-4 litters (W). Measurements of the distance from MCC nuclei to stroma in the INF/AMP region of dams with 3-4 litters or 7-8 litters (Y). Each point represents a measurement/cell. The continuous line represents the median, while the dotted lines show the quartiles. **** = p < 0.0001. (Z, a) 13mo mice that have had 7-8 litters in their lifetime had larger and less atrophic ovaries (a), as compared to those with 3-4 litters (Z). OvA: ovarian artery, UtA: uterine artery, Ov: ovary, Ovi: oviduct, Ut: uterus.

Since aging-associated decline in female fertility is caused by ovarian aging/oocyte depletion, we wondered if MCC vacuolation contributed to this decline in fertility by hindering OCC movement into the oviduct. Regardless of cytoplasmic vacuoles in MCCs, superovulation of 13mo mice resulted in the successful entry of OCCs into the oviduct (Fig. 7V), indicating that vacuolation did not affect proper INF/AMP functions mediated by MCC beating (2). Indeed, INF/AMP MCCs in aged 13mo mice were still beating (Suppl. Video 1), suggesting continued ATP consumption of MCCs in aged oviducts.

Pregnancy modulates blood circulation around the female reproductive tract to support fetal growth. It is shown that the uterine artery contributes 68%, and the uterine branch of the ovarian artery contributes 32% to the placenta/fetuses (36), indicating that the ovarian artery also supports the continuation of pregnancy. We wondered whether gravidity, defined as number of pregnancies, would impact INF/AMP MCC vacuolation. Interestingly, INF/AMP MCCs in 13mo mice that have given birth to 7-8 litters in their lifetime (last litter at age 8-9mo/4-5 months prior to euthanasia) presented with decreased frequency of vacuolation and apical displacement of nucleus, relative to 13mo mice that have given birth to 3-4 litters in their lifetime (last litter at age 5-6mo/7-8 months prior to euthanasia) (Fig. 7W-Y). We also found that 13mo mice with 7-8 litters had larger and less atrophic ovaries, as compared to mice with 3-4 litters (Fig. 7Z, a). As expected, MCC vacuolation was restricted to the INF/AMP epithelium and not noted in the other regions of the oviduct (Suppl. Fig. 7). Taken together, we propose that changes in blood circulation via the ovarian artery contribute to ovarian and distal oviduct aging. Reduced blood circulation results in a constant, mildly stressed microenvironment within the INF/AMP region leading to MCC vacuolation.

## DISCUSSION

In this study, we show that the INF/AMP MCCs uniquely present with aging-associated cytoplasmic vacuolation and nuclear anomaly. We initially suspected that repetitive exposure to inflammatory factors or ROS released during ovulation (19) could be part of the cause. However, ovariectomy did not affect the timeline of MCC vacuolation in the INF/AMP epithelium. Additionally, the MCC vacuolation phenotype was restricted to the INF/AMP epithelium and not observed in the AIJ epithelium despite the same proximity to the ovary in a single transverse section. We concluded that MCC vacuolation in the INF/AMP epithelium was not associated with proximity to the ovary but was due to a distinct cellular subtype and microenvironmental changes, suggesting intrinsic vulnerability.

MCCs in the INF/AMP epithelium are constantly beating regardless of the female hormonal cycles and ovulation. Although changes in cilia beat frequencies (CBF) are reported during the different hormonal stages, it does not impact OCC transport along the AMP (4). We observed that MCCs in the aged INF/AMP continued to beat and transport OCCs into the AMP despite MCC vacuolation. In *in vitro* organotypic slice cultures, inhibition of cellular respiration terminated cilia beating (data not shown) and resulted in INF/AMP MCC protrusion/death within 24 hours of treatment, with no noticeable changes in SCs or AIJ/ISM MCCs. This is consistent with a recent study showing that lung MCCs and their progenitors respond differently to mitochondrial dysfunction (27). Taken together, this suggests that INF/AMP MCCs rely heavily on oxidative phosphorylation to fulfill high ATP usage required for constant cilia beating.

The most ATP-consuming biochemical events in adult cells are 1) Na+, K+ ATPase activity responsible for the establishment and maintenance of ion homeostasis, membrane potential, active transport of substrates, and 2) protein turnover (29, 37). However, constant MCC beating in the INF/AMP epithelium continued even after the cessation of ovarian functions and MCC vacuolation, suggesting that these cells prioritize cilia function/motility. Acute ATP depletion induced by combined inhibition of glycolysis and cellular respiration in MDCK cells results in the internalization of Na+, K+ ATPase and E-cadherin (26). Thus, the cytoplasmic vacuoles lined with Na+, K+ ATPase α1 subunit (ATP1A1) observed in INF/AMP MCCs of aged mice could be indicative of cellular adaptation to ATP shortage.

MCC vacuolation was induced in hypoxia (3% O_2_) or high dose HU treatment in *in vitro* slice cultures, but not by ER stress inducers or inhibition of protein synthesis, glycolysis, hexosamine biosynthesis or pentose phosphate pathway. Acute hypoxia promotes Na+, K+ ATPase internalization by generating mitochondrial ROS (38), while chronic hypoxia results in ROS elevation and diminished ATP demand (29). In line with this, INF/AMP MCCs in 18mo mice showed downregulation of gene sets associated with cilia function/motility, reflecting decreased ATP demand, and upregulation of gene sets associated with ROS metabolism/catabolism, suggesting that the MCCs are adapting to environmental stress.

HU treatment, known to inhibit DNA synthesis, resulted in MCC vacuolation. HU predominantly affects the S-phase of the cell cycle, inducing dNTP depletion and replication fork stalling in cycling cells (31) by inhibiting ribonucleotide reductase (RNR). Hypoxia also induces dNTP depletion and replication stress in cycling cells (39), as oxygen is an essential co-factor for RNRs (40). Based on our finding of phospho-H2AX puncta in INF/AMP MCCs throughout adulthood and aging, it is interesting to speculate that vacuolation may be a response to replicative stress, despite MCCs being terminally differentiated, non-cycling cells. Interestingly, prolonged HU treatment results in formation of heterogeneous numbers of centrosomes in cycling Chinese Hamster Ovary (CHO) cells, suggesting that HU may countermand the suppression of centriole/centrosome formation (41). Thus, in non-dividing MCCs with numerous centrioles, another possibility is that prolonged, high-dose HU treatment deregulates centriole numbers.

Although MCCs from all three regions of the oviduct were sensitive to hypoxia and presented vacuolation phenotypes *in vitro*, the MCC vacuolation phenotype *in vivo* was restricted to the INF/AMP epithelium. This suggests not only intrinsic vulnerability but also unique microenvironmental differences in the INF/AMP epithelium.

Three arteries feed into the female reproductive tract: ovarian, uterine and cervical arteries (32). We found that the ovarian artery feeds into the ovary and INF/AMP region, while the uterine artery supports the proximal oviduct. Interestingly, in some Mayer-Rokitansky-Küster-Hauser (MRKH) syndrome patients, the distal end of Fallopian tube is intact despite the lack of the uterus (42), suggesting that its circulation is independent of the uterus. Branches of the ovarian artery connect into the distal region, while the uterine artery feeds into the proximal Fallopian tube (43). Thus, blood circulation patterns in mice and humans are conserved despite apparent anatomical differences between mouse and human female reproductive tracts.

Our data showed blood circulation to the INF/AMP region relied on the ovarian artery, which was tightly linked to ovarian function. In aging, we found reduced circulation to the ovary and the INF/AMP. We propose that this creates a chronic, mild low oxygen and low nutrient condition in the INF/AMP region, reflected by MCC vacuolation.

Finally, the distal fimbria of the human Fallopian tube is recognized the site of origin of High Grade Serous Ovarian Carcinomas (HGSCs) (10). It is interesting to speculate that low blood circulation to the distal Fallopian tube via the ovarian artery in postmenopausal individuals could lead to a constant, mild environmental stress that facilitates the initiation and progression of HGSCs. In one human study, while variations exist between individuals and hormonal conditions, changes in the ovarian/uterine artery calibers are observed in FTs of cycling and postmenopausal women, suggesting that FTs from cycling humans receive blood supply mainly from the ovarian artery and FTs from postmenopausal humans predominantly receive blood supply from the uterine artery (44). Further, we also showed that parity was directly proportional to decreased severity of MCC nucleus displacement and vacuolation, suggesting that pregnancy provided a protective effect. Likewise, the *P53* signature, considered a latent HGSC precursor, is significantly associated with lower parity (45), and thus with increased number of ovulations. While cytoplasmic vacuolation occurred independent of estrous cycling and/or ovulation, the severity of vacuolation decreased with higher gravidity/parity. Pregnancy involves increased blood flow and oxygenation of the reproductive tract (36), including the distal oviduct and ovary, which also prevents shrinkage and decreased atrophy of ovaries in mice with higher gravidity.

Taken together, we propose that the INF/AMP epithelium has unique intrinsic properties and circulatory environment, as compared to the rest of the oviduct. Changes in blood circulation were associated with cessation of ovarian function and resulted in environmental stress that was reflected in INF/AMP-specific MCC vacuolation.

## MATERIALS AND METHODS

### Sex as a biological variable

Only female animals were used in this study on organs of the female reproductive tract.

### Animals

Our study exclusively examined female mice because the organ is only relevant in females. All animal work was performed in accordance with institutional guidelines and was approved by the Faculty of Medicine Animal Care Committee (AUP #7843); it was undertaken at the Goodman Cancer Research Institute animal facility. Aging phenotypes were confirmed in *Fltp-H2B-Venus* (46) mice that were a kind gift from Dr. Heiko Lickert and Tdtomato flox/flox mice (Ai14) acquired from JAX (#007914). Wild-type B6N mice acquired from McGill’s Integrated Core of Animal Modeling (MICAM) were used for ovariectomy surgeries (N=3-4 for each time-point), and results were confirmed in *Fltp-H2B-Venus* mice (N=2-3 for each time-point). 13mo retired B6N dams with 3-4 (N=3) or 7-8 litters (N=3) were acquired from MICAM, where trio breeding is the standard. Where required, estrous stages were analyzed using vaginal smears stained with crystal violet (47).

### 3D confocal imaging

Oviducts were collected and straightened by removing the mesosalpinx. The straightened oviduct was fixed with DMSO (Dimethyl sulfoxide, Sigma Aldrich D8418): methanol (BioShop Canada Inc., MET302.1) in the ratio 1:4 and placed at −20°C overnight. The antibody staining protocol was as described in (48). Blocking was done overnight in solution containing 1% Triton X-100 (Sigma-Aldrich, T8787), 2% powdered milk, and 1X phosphate-buffered saline (PBS, BioShop Canada Inc., PBS405), henceforth referred to as PBSMT. Primary and secondary antibody staining was performed in PBSMT for 5 and 2 days, respectively, at 4° on a shaker. After six 30-minute-long PBSMT washes, oviducts were transferred successively to methanol:PBT (1:1, PBT: 1× PBS and 1% Triton X-100), 100% methanol, 3% H2O2 (BioShop Canada Inc., HYP001.1), and 100% methanol prior to benzyl alcohol/benzyl benzoate (BABB) clearing. BABB-cleared samples were placed on a #1.5 coverslip (Fisher Scientific, 12-545F) with 10–15 μl BABB prior to imaging using the 10X objective (0.30 numerical aperture/N. A) on LSM 800 or 710 (Zeiss). Section interval for 3D confocal imaging was 4.32 μm. Usually, 40–110 optical sections were taken.

### Bilateral and unilateral ovariectomy surgeries

Surgery preparation and female reproductive tract exposure was as described previously (49). Mice were anesthetised with isoflurane (McGill’s Comparative Medicine and Animal Resources Centre/CMARC pharmacy) and injected with 20mg/kg subcutaneous Carprofen (CMARC pharmacy) prior to fur removal and surgery. A dorsal midline skin incision (∼1 cm) was created, followed by an incision in the body wall located directly above the ovarian fat pad. This ovarian fat pad was pulled out using a pair of curved forceps (Fisher Scientific, NC0696845), exposing the upper female reproductive tract (ovary, oviduct, uterus). The tract was placed on a sterile gauze soaked in saline and a bulldog clamp (Fisher Scientific, 50-822-230) was secured on the fat pad to keep the tract exposed. Under the microscope (SteREO Discovery.V8, Zeiss), the ovarian bursa was cut open to expose and cut out the ovary using a pair of sterile microdissection scissors (Fisher Scientific, 089531B). Following ovary removal, body wall and skin incisions were sutured close using absorbable vicryl sutures (Fisher Scientific, 50-118-0847). Mice were administered Carprofen for 3 days and monitored for 10 days.

### Immunofluorescence and immunohistochemistry

Uncoiled or coiled oviducts/reproductive tracts were fixed with 4% paraformaldehyde (PFA, Polysciences) for 30 minutes, followed by three washes with 1X PBS. For cryosectioning, fixed samples were calibrated in 15% and 30% sucrose either at 4 degrees overnight prior to OCT (Fisher HealthCare, #23-730-571) embedding and sectioning at 5-7-or 10-micron thickness. For immunofluorescence (IF), sections were brought to room temperature and washed with 1X PBS, 0.1% Triton X-100, and 0.5% Triton X-100. Slides were incubated in permeabilization and blocking solution (0.1% Triton X-100 and 1% FBS in 1X PBS) for 2 hours at room temperature. Primary antibody staining was performed overnight at 4°C in permeabilization and blocking solution. Primary antibodies were washed off with multiple washes of 1X PBS and 0.1% Tween (Sigma Aldrich, P9416), followed by secondary antibody staining in 1X PBS overnight at 4°C. After the secondary antibodies were washed off, 1-2 drops of homemade mounting media were pipetted onto the sections and #1.5 coverslips were placed over the stained sections. Coverslips were sealed to the slides using nail polish. IF sections were imaged with 20x (N.A 0.8) or 63x (N.A 1.4) objectives, on LSM710 or LSM800 confocal microscopes (Zeiss). Homemade mounting media was made of 1-part 10X PBS, 9 parts glycerol (BioShop, GLY001), and slowly stirred in 0.1 parts of 20% w/v propyl gallate (Sigma Aldrich, P3130; dissolved in DMSO). Paraffin processing, Haematoxylin & Eosin (H&E), immunohistochemistry (IHC), and digital slide scanning (NanoZoomer S210, Hamamatsu) were performed by the Goodman Cancer Research Institute Histology Innovation Platform. Embedded blocks were sectioned on the microtome at 4– 6-micron thickness. For IF on paraffin sections, deparaffinization involved heating at 50°C, two incubations of 5 minutes each in Histo Clear (National Diagnostics, HS-202) and 3-minute-long incubations in successive ethanol solutions (100%, 90%, 70%, 50%, 0% in water). Antigen retrieval was performed with homemade 1X buffer (20 mM Tris, 1mM EDTA, 0.05% Tween-20 at pH 9.0) for 7 minutes in a pressure cooker. Primary and secondary antibody staining was performed as stated above.

150µm thick organotypic slice cultures were transferred to 4-well plates and fixed in 4% PFA for 30 minutes at room temperature, on a shaker. PFA was replaced by 1X PBS thrice, each wash with shaking for 10 minutes at room temperature. 1X PBS was replaced by permeabilization and blocking solution and incubated overnight at 4°C on a shaker. Primary antibody staining was performed in permeabilization and blocking solution overnight at 4°C on a shaker, followed by multiple 1X PBS and 0.1% Tween washes, secondary antibody staining in 1X PBS on day 3, and multiple washes. Slices were transferred to a #1.5 coverslip with a 120µm spacer stuck on the surface. Any solution remaining on the coverslip was replaced by a drop of homemade mounting media, followed by a second #1.5 coverslip on top, such that the slices were sandwiched between two coverslips. Stained slices were imaged with a 20x objective (0.8 N.A) on LSM710 or LSM800 confocal microscopes.

### Antibodies

Primary antibodies (1/250 dilution, unless specified): anti-SOX17 (R&D Systems, AF1924), anti-PAX8 (Proteintech, 10336-1-AP), anti-FOXJ1 (Abcam, ab235445), anti-mcherry (Abcam, ab213511/Invitrogen, #M11217), anti-GFP (Abcam, ab13970), anti-ATP1A1 (a2F, DSHB), anti-LaminB1 (Cell Signaling, 13435S), anti-ᵞH2AX (Cell Signalling, 9718T), anti-P16 (Cell Signalling, 92803T), anti-P21 (Cell Signalling, 2947T).

Secondary antibodies (1/450 dilution): Alexa Fluor (AF) anti-rabbit 555 (Invitrogen, A31572), AF anti-rabbit 649 (Invitrogen, A21245), AF anti-rabbit 488 (Invitrogen, A21206), AF anti-mouse 649 (Invitrogen, A32787), AF anti-mouse 488 (Invitrogen, A21202), AF anti-goat 568 (Invitrogen, A11057), AF anti-rat 488 (Invitrogen, A21208), anti-chicken 488 (Sigma, SAB4600031), DAPI (4′, 6-diamidino-2-phenylindole, Thermo Fisher, 62248), Hoescht 33342 (Thermo Fisher, 62249), AF 488 phalloidin (Lifetech, A12379), and AF 635 phalloidin (Lifetech, A34054).

### Isolation of INF/AMP cells for single cell RNA sequencing

Oviducts from a single 3mo mouse in estrus (Estrus_INFAMP) or three 18mo mice (Aged_INFAMP) were straightened out in 1X PBS. The INF/AMP region was separated from the rest of the tube by slicing in between turns 1 and 2. INF/AMP regions were moved into media containing 1X PBS, 100 IU/ml of Penicillin & 100ug/ml of Streptomycin (Gibco, 15140122), and 10% fetal bovine serum (FBS, Wisent Bioproducts, 090-150). These pieces of tissue were transferred onto empty plates with a drop of media containing 5mg/ml collagenase B (Sigma-Aldrich, COLLB-RO) and 5U/100ul DNase I (Thermo Scientific, EN0521) and chopped into smaller pieces with a No. 21 blade (Paragon, P307). This solution with small tissue pieces was transferred to a 1.5ml tube containing 100µl of the above-mentioned dissociation solution and incubated for 35-40 minutes at 37°C. This was then spun down at 1,500rpm for 5 minutes, resuspended in 500µl 1X PBS, and spun down again to resuspend in 50µl of warmed 0.25% Trypsin-EDTA (Gibco, Cat# 25200056) followed by a 7-minute incubation at 37°C. Trypsinization was halted with media, and mechanical dissociation was performed by passing through a 27 ½ gauge insulin needle (BD, #324704). The resulting single cell suspension was passed through a 40µm cell strainer (Corning, 352340), centrifuged at 1,500rpm for 7 minutes and resuspended in 200µl 1X PBS.

### Single Cell RNA sequencing (scRNASeq)

Quality control and scRNASeq was performed by the McGill Genome Centre. For initial quality control, an aliquot was taken from the cell suspension and incubated in a working concentration of 2mM calcein-AM and 4mM Ethidium-Homodimer1 (Thermofisher L3224). After 10 minutes of incubation at room temperature, sample viability, concentration, segregation, size and absence of large debris were verified by loading stained cell suspension onto hemocytometer (Incyto DHC-N01-5) and imaged on bright field, GFP (for Calcein-AM) and RFP (for Ethidium homodimer-1) channels using a EVOS FL Auto Fluorescent microscope (ThermoFisher). Samples with a viability of 70% or more proceeded to scRNASeq. Single cell gene expression data was generated according to Chromium Single Cell User Guide (v3.1 Chemistry, 10X genomics). Briefly, cells were suspended into Reverse Transcription (RT) Master Mix (10X genomics) then pipetted into Well-1 of a Chip ‘‘G’’ (10X genomics), followed by Gel Beads (10X genomics) and Partition oil (10X genomics). The chip assembly was run on a Chromium Controller (10X genomics) which generated Gel Bead-In-EMulsions (GEMs). Following RT protocol on a thermocycler (Biorad T100), emulsion beads were broken, and quality (size distribution) and concentration of cDNA was assessed using LabChip (Perkin Elmer 760517, CLS760672). Barcoded cDNAs were pooled for amplification, and the final PCR product (or sequence ready library) was purified, and quality controlled using LabChip. Finally, the libraries were sequenced on 1 lane of MGI G400, DNBSEQ-G400 sequencer (MGI Tech Co. Ltd).

### scRNASeq analyses

Raw sequencing data for each sample was converted to matrices of expression counts using the Cell Ranger software provided by 10X genomics (version 2.0.2). Briefly, raw BCL files were demultiplexed into paired-end, gzip-compressed FASTQ files for each channel using Cell Ranger’s mkfastq. Using Cell Ranger’s count, reads were aligned to the mouse reference transcriptome (mm10), and transcript counts quantified for each annotated gene within every cell. The resulting UMI count matrix (genes 3 cells) were then provided as input to Seurat suite (version 2.3.4) (50). Cells were first filtered to remove those that contain less than 200 genes detected and those in which > 10% of the transcript counts were derived from mitochondrial-encoded genes. Estrus_INFAMP and Aged_INFAMP samples were merged into one 10x combined object using canonical correlation analysis (CCA), followed by scaling data (ScaleData function) and finding variable genes (FindVariableGenes function). CCA subspaces were aligned using CCA dimensions 1-15, which was followed by clustering (FindClusters function) and integrated UMAP visualization for all cells. Cluster-specific gene markers were identified using Seurat’s FindMarkers with cutoffs log2FC > 1 and P-adjusted value < 0.05. Gene Set Enrichment Analysis (GSEA) was performed per cluster with cutoff P-adj < 0.05 using gene sets of Gene Ontology (GO) biological processes.

### Live imaging cilia beating

INF regions of the oviduct were cut off and sliced radially in warm 1X PBS. These pieces were gently transferred to a #1.5 coverslip (VWR, 48393-251) with a 120µm spacer (Invitrogen, S24737) stuck on the surface, and any remaining PBS was replaced with a ∼10µl of warmed media containing Advanced DMEM/F12 (Gibco, 12634010), 2mM GlutaMax (Gibco, #35050061), 5% FBS, and 50U/ml penicillin/streptomycin. A second #1.5 coverslip was placed on top to sandwich the tissue pieces, and they were imaged within 30 minutes using a 20X objective (0.75 N.A) on a widefield microscope system (Nikon), built around a Nikon TI2-E stand, equipped with Nikon‘s Perfect Focus System, a LED light source (SpectraX, Lumencor) and sCMOS cameras (Orca Fusion-BT, Hamamatsu) enclosed by a custom-built environmental chamber (Digital Pixel), and controlled using Nikon NIS Elements AR software. 2.5-second-long, >400 fps, brightfield, 512 x 512 ROI videos of tissue edges covered with MCCs were imaged at 37°C.

### Superovulation

13mo mice were administered 100µl of CARD HyperOva (Cosmo Bio Ltd., KYD-010-EX) intraperitoneally at 5pm. 48 hours later, 7.5 IU of human Chorionic Gonadotropin (hCG; Sigma, CG-10) was administered intraperitoneally. Mice were euthanized 15-17 hours after hCG injection, and tracts were isolated for paraffin processing.

### Organotypic slice culture

Reproductive tracts were dissected out of 2-3-month-old mice and dropped into a petri dish containing ice-cold 1X PBS to reduce contractions. Ovaries were removed, and oviducts were straightened out by removing the mesosalpinx. ∼1ml of warmed 4% low melting agarose (LMA, IBI Scientific, IB70051) was pipetted into a cryomold (Fisherbrand, 22363553). The straightened oviduct was embedded in the LMA and placed on ice to set. A portion of the uterus was kept intact to stabilize the tract in the LMA while sectioning. The solidified LMA block was superglued (Krazy Glue) onto the metal specimen holder such that the cut side of the uterus faced the specimen holder, and the distal tip of the uncoiled oviduct faced the ceiling. After the glue dried, the specimen holder was placed in the buffer tray containing ice cold 1X PBS, surrounded by ice in the ice bath tray. The block was sectioned into 150µm thick, transverse slices at a speed and amplitude of 0.8mm/s and 1.60mm respectively, using a vibratome (Leica). Slices were picked up using a brush and placed in a well of a 24-well-plate (Fisher Scientific, FB012929) containing 250µl of ice-cold media. Tissue slices embedded in agarose were transferred to a 4-(Fisher Scientific, Cat# FB012926) or 25cm^2^ dish (Fisher Scientific, 08-757-500) and 500µl/8ml of fresh, ice-cold media was added to each well/dish. Media contained Advanced DMEM/F12, 2mM GlutaMax, 5% FBS, and 50U/ml penicillin/streptomycin.

25cm^2^ dishes were used for experiments comparing normoxia to 3% hypoxia, while 4-well or 24-well plates were used for all other treatments. Organotypic slice cultures in 25cm^2^ dishes were placed in normoxia or hypoxia on the day of sectioning. Hypoxia (3.5% O_2_, 5% CO_2_, 91.5% N_2_) was maintained in a separate, humidified CO_2_ incubator with O_2_ control (Heracell 150i, ThermoFisher). Organotypic slice cultures in 4-well plates were placed in a humidified CO_2_ incubator for 24 hours prior to inhibitor treatments. Inhibitors/treatments used were 200µM/4mM hydroxyurea (HU, Sigma Aldrich, H8627-5G), 10µg/ml cycloheximide (CHX, Cayman Chemical Company, 14126), 2µM 6-Aminonicotinamide (6-AN, Sigma, A68203-1G), 1µM YZ-9 (Cayman Chemical Company, 15352), 5µM tunicamycin (TUN, Sigma Aldrich, T7765-10MG), 1µM shikonin (Cayman Chemical Company, 14751), 0.8µM ST045849 (Sigma Aldrich, SML2702-5MG), 10µM Antimycin-A (Sigma Aldrich, A8674) and/or 10mM 2-deoxyglucose (2-DG, Sigma Aldrich, D6134).

### Cardiac perfusion for blood vessel visualization

Mice were anesthetized with isoflurane, and cardiac chamber was opened to expose the heart. A catheter was inserted into the left ventricle, the right ventricle was cut open, and 30ml 0.85% saline was injected to flush the arteries. After flushing, 4ml of 0.25% CellMask (Invitrogen, C10045), diluted in 0.85% saline, was injected into the left ventricle. The female reproductive tract was dissected and imaged under a stereomicroscope (SteREO Lumar. V12, Zeiss).

### Image analysis and statistics

Stained sections were visualized using FIJI, Zen (Zeiss) or ImageScope x64 (Aperio). Apical displacement of nuclei was measured manually on FIJI by drawing a line from the base of nucleus to the basal lamina/stroma as parallel to the lateral domain as possible, using either 63x IF images taken on a confocal microscope or 40x H&E-stained digital slides that were converted to TIFF format using ndpitools and ndpi2tiff plugins. Cellular/epithelial monolayer height was measured manually on FIJI by drawing a line from base to apical side of MCCs. Measurements were performed from base to base for each mucosal fold to include cells at the base, side and tip of each mucosal fold. Apical displacement and cell height measurements were compared between mice of the same genotype and sample preparation.

Cell counts were performed on 20x confocal images using the Cell Counter plugin. Although FLTP-H2B-Venus and FOXJ1 were used interchangeably as MCC markers, cell counts were compared only between samples stained with the same marker. Number of MCCs in organotypic slice cultures were first counted on a single z-stack, and then vacuolated MCCs were identified by inspecting the neighboring stacks. GraphPad Prism was used to perform unpaired, two-tailed t-test and create graphs/plots.

## Authors’ contributions

KH executed and analyzed most experiments. NY performed the ovariectomy surgeries and provided aged mice with multiple litters. ASP performed bioinformatic analyses. SZ, AFT, VL, MT, KT, and MJF performed some sectioning and/or staining. YY and WP performed the blood vessel visualization experiments. KH, WP, MJF, and YY edited the manuscript. KH and YY conceived the project and wrote the manuscript.

## Acknowledgements

This work was supported by Canadian Cancer Society (CCS) Innovation Grant (Haladner Memorial Foundation #704793), CCS innovation to Impact (i2I) Grant (#706320), and Cancer Research Society Operation Grant (#23237). KH was supported by Charlotte and Leo Karassik Foundation Oncology Studentship, Fonds de recherche du Québec – Santé (FRQS) doctoral fellowship, DKG World Fellowship 2021, Donner Foundation, Hugh E. Burke, Centre for Research in Reproduction & Development (CRRD), Alexander McFee, and Rolande & Marcel Gosselin graduate studentships. WP was supported by the CRRD trainee fellowship. MJF was supported by Canderel, CRRD, and FRQS postdoc fellowships. KT was supported by McGill Integrated Cancer Research Training (MICRTP) and Canderel graduate studentships.

The authors would like to thank Dr. Arnold Hayer, Dr. Alain Nepveu, and their lab members for sharing equipment and help with cilia live-imaging and hypoxia experiments. We would also like to thank Dr. Yu Chang Wang, Mr. Haig Hugo Vrej Djambazian and Dr. Jiannis Ragoussis (McGill Genome Centre) for help with the scRNASeq. As well, we thank the Rosalind and Morris Goodman Cancer Research Institute Histology Facility, McGill’s Advanced Bioimaging Facility (ABIF), McGill Genome Centre, and the McGill Integrated Core of Animal Modeling (MICAM) for technical support.

