## Supplementary Figures for "Aging-associated vacuolation of multi-ciliated cells in the distal mouse oviduct reflects unique cell identity and luminal microenvironment"

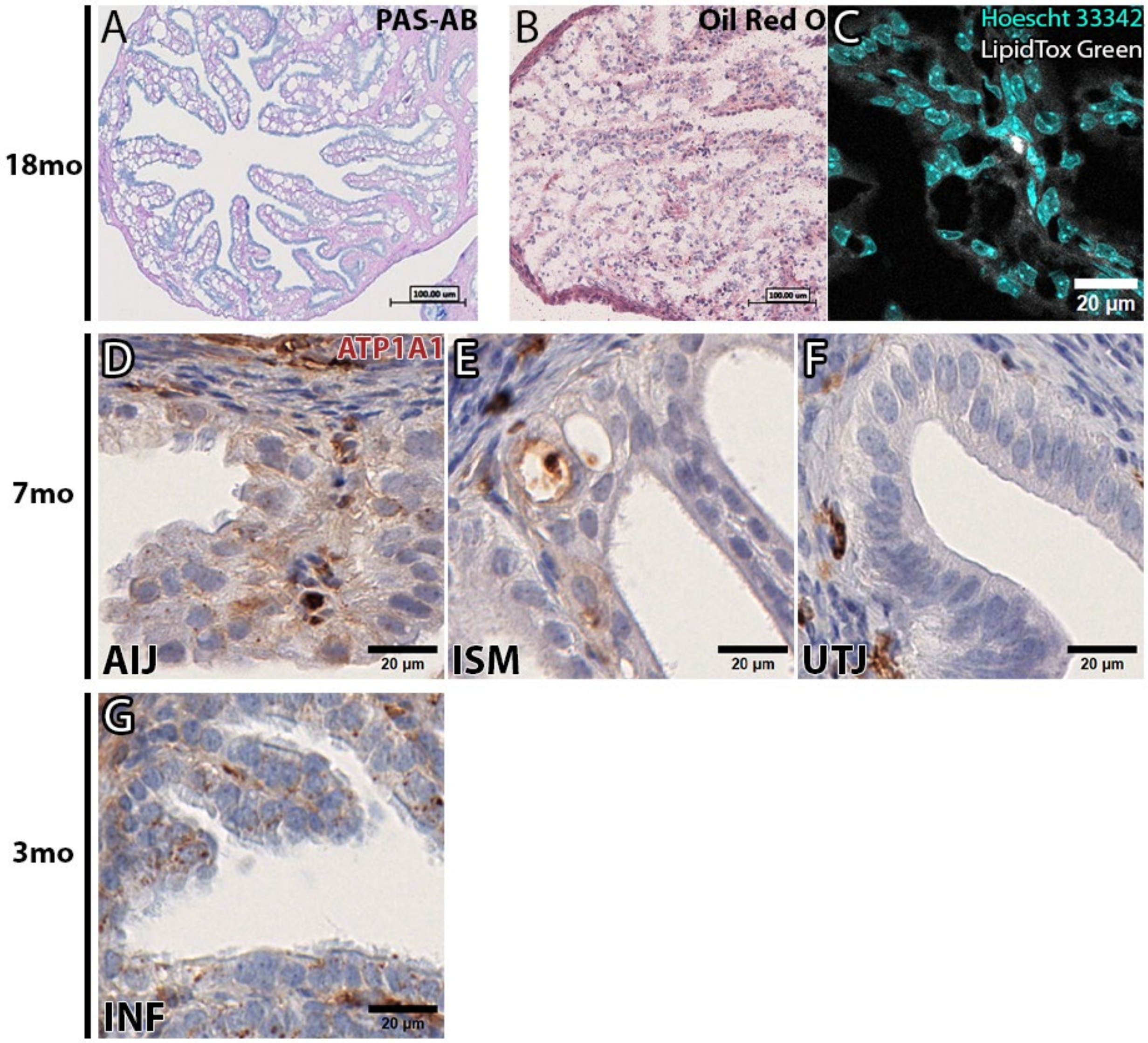


**Suppl. Fig. 1: Aging-associated cytoplasmic vacuoles in INF/AMP MCCs do not contain glycogen or lipids, and no ATP1A1 localization was noted in the proximal oviduct.** (A) Periodic acid-Schiff/PAS staining of the 18mo INF/AMP region. Cytoplasmic vacuoles were negative for PAS, indicating that vacuoles do not contain polysaccharides like glycogen or mucus-like substances. (B, C) Lipid staining of the 18mo INF/AMP region. Cytoplasmic vacuoles were negative for Oil Red O (B) and LipidTox Green (C), indicating that vacuoles do not contain lipid.

(D-F) ATP1A1 IHC staining in the AIJ (D), ISM (E) and UTJ (F) regions isolated from 7mo mice, showing no specific localization to lateral surfaces or in the cytoplasm. (G) ATP1A1 IHC staining in the INF/AMP region isolated from 3mo mice, showing occasional punctate staining.

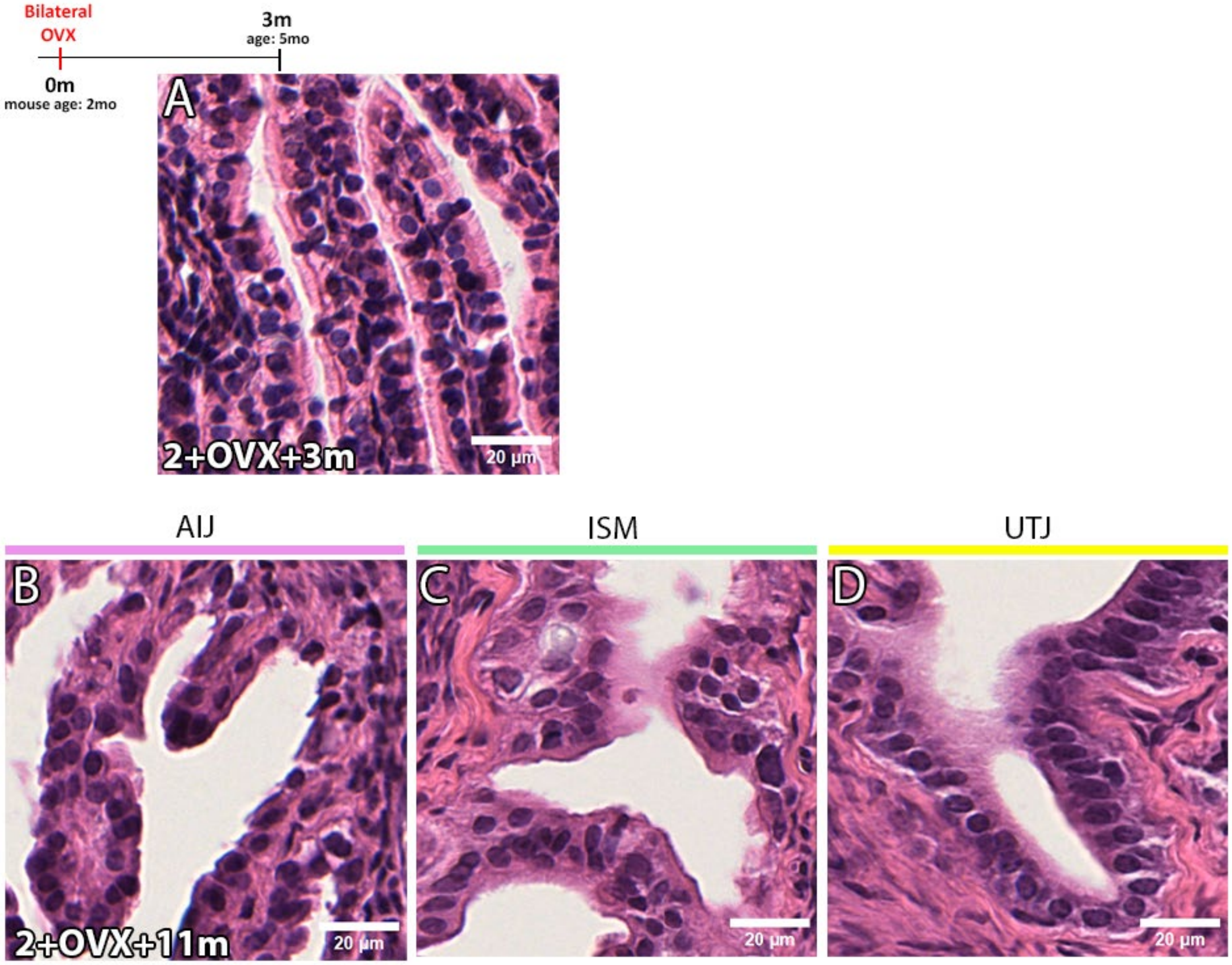


**Suppl. Fig. 2: No discernible vacuoles in the INF/AMP region of mice 3 months following OVX and in the AIJ, ISM, and UTJ regions of mice 11 months post OVX.** (A) H&E staining of the INF/AMP region in oviducts from mice 3 months post bilateral OVX. No discernible cytoplasmic vacuoles were noted. (B-D) H&E staining of the AIJ (B), ISM (C) and UTJ (D) region in oviducts from mice 11 months post bilateral OVX. No discernible cytoplasmic vacuoles were noted.

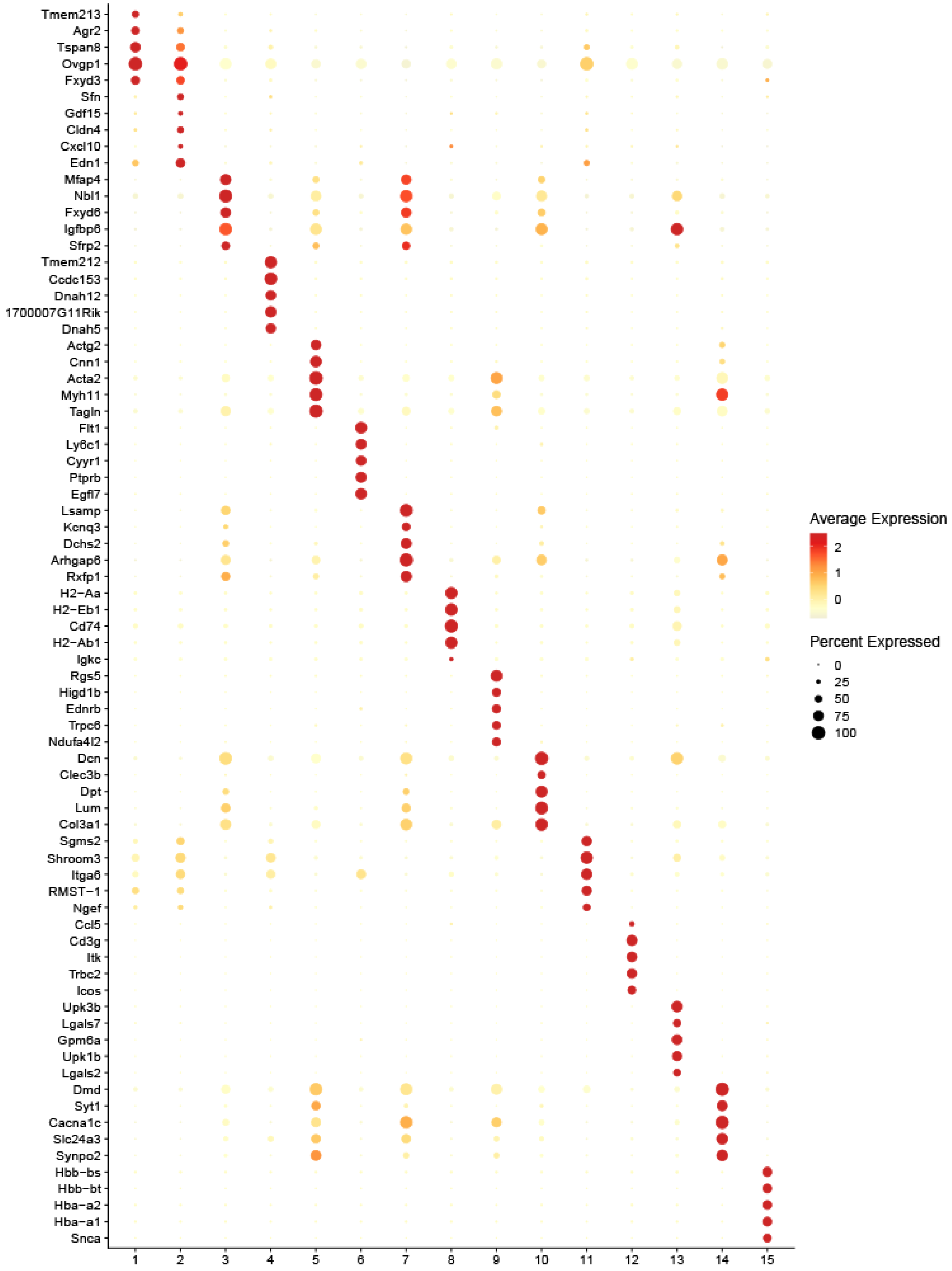

**Suppl. Fig. 3:** Dotplot of top 5 markers in each cluster.

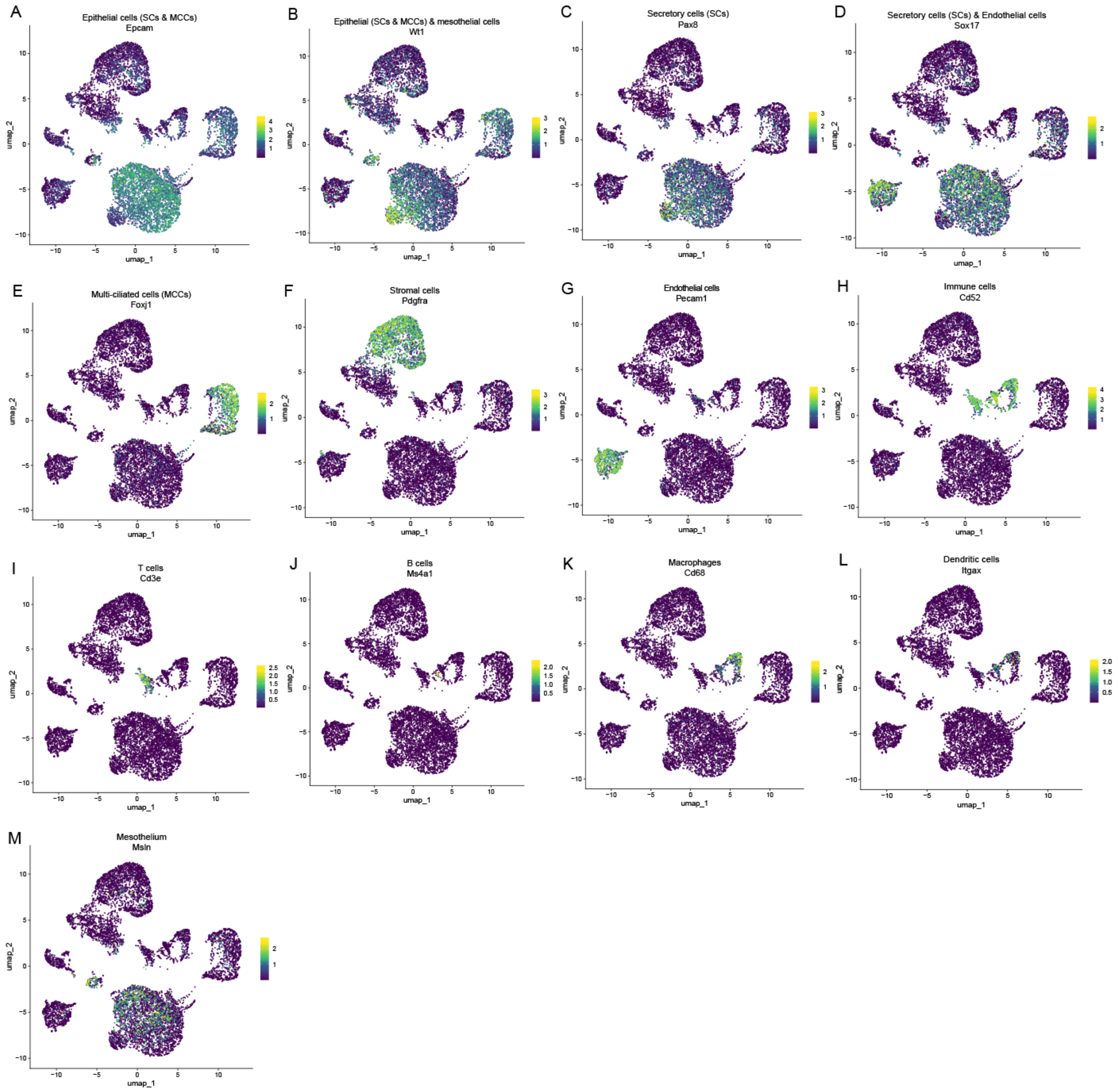


**Suppl. Fig. 4: Identification of isolated cell populations using gene expression of known markers.** (A-M) Gene expression of known markers that were not included in the dotplot, such as Epcam (A), Wt1 (B), Pax8 (C), Sox17 (D), Foxj1 (E), Pdgfra (F), Pecam1 (G), Cd52 (H), Cd3e (I), Cd20 (J), Cd68 (K), Cd11c (L), Msln (M).

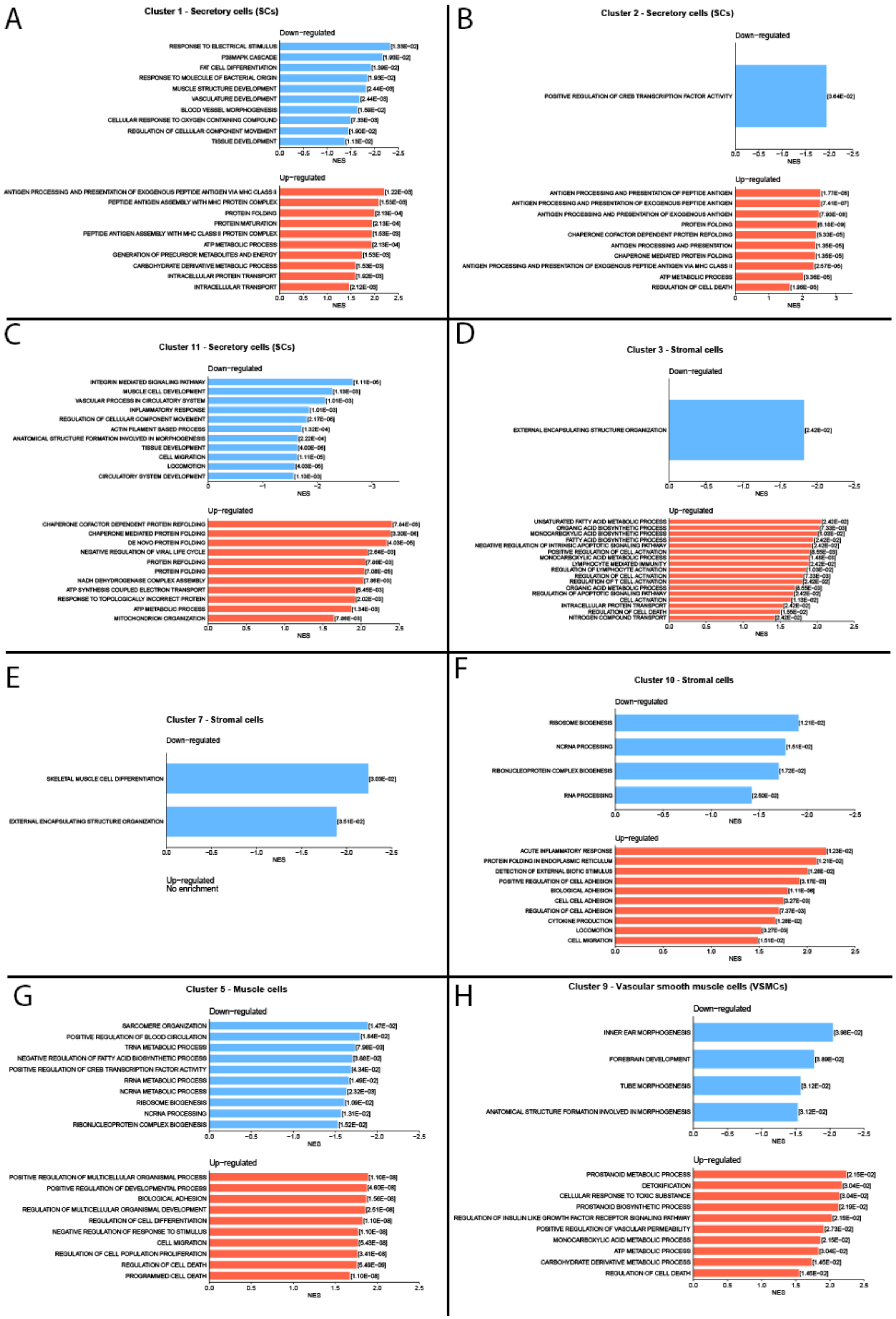

A
Cluster 1 - Secretory cells (SCs)
B
Cluster 2 - Secretory cells (SCs)
C
Cluster 11 - Secretory cells (SCs)
D
Cluster 3 - Stromal cells
E
Cluster 7 - Stromal cells
F
Cluster 10 - Stromal cells
G
Cluster 5 - Muscle cells
H
Cluster 9 - Vascular smooth muscle cells (VSMCs)
Down-regulated
Up-regulated
NES
Up-regulated
No enrichment

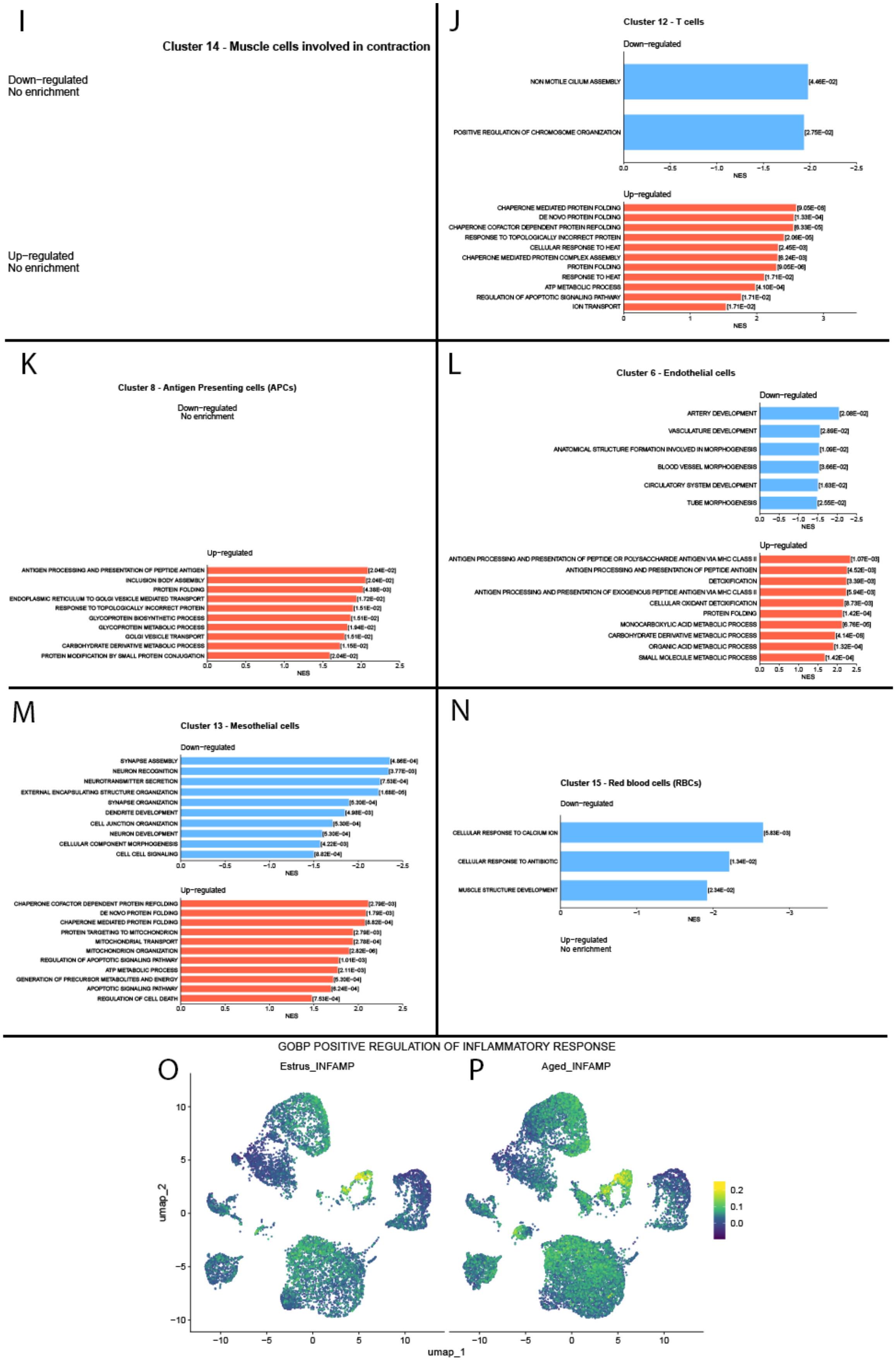
I
Cluster 14 - Muscle cells involved in contraction
Down−regulated
No enrichment
Up−regulated
No enrichment
J
Cluster 12 - T cells
Down−regulated
NON MOTILE CILIUM ASSEMBLY
POSITIVE REGULATION OF CHROMOSOME ORGANIZATION
Up−regulated
CHAPERONE MEDIATED PROTEIN FOLDING
DE NOVO PROTEIN FOLDING
CHAPERONE COFACTOR DEPENDENT PROTEIN REFOLDING
RESPONSE TO TOPOLOGICALLY INCORRECT PROTEIN
CELLULAR RESPONSE TO HEAT
CHAPERONE MEDIATED PROTEIN COMPLEX ASSEMBLY
PROTEIN FOLDING
RESPONSE TO HEAT
ATP METABOLIC PROCESS
REGULATION OF APOPTOTIC SIGNALING PATHWAY
ION TRANSPORT
NES
K
Cluster 8 - Antigen Presenting cells (APCs)
Down−regulated
No enrichment
Up−regulated
ANTIGEN PROCESSING AND PRESENTATION OF PEPTIDE ANTIGEN
INCLUSION BODY ASSEMBLY
PROTEIN FOLDING
ENDOPLASMIC RETICULUM TO GOLGI VESICLE MEDIATED TRANSPORT
RESPONSE TO TOPOLOGICALLY INCORRECT PROTEIN
GLYCOPROTEIN BIOSYNTHETIC PROCESS
GLYCOPROTEIN METABOLIC PROCESS
GOLGI VESICLE TRANSPORT
CARBOHYDRATE DERIVATIVE METABOLIC PROCESS
PROTEIN MODIFICATION BY SMALL PROTEIN CONJUGATION
NES
L
Cluster 6 - Endothelial cells
Down−regulated
ARTERY DEVELOPMENT
VASCULATURE DEVELOPMENT
ANATOMICAL STRUCTURE FORMATION INVOLVED IN MORPHOGENESIS
BLOOD VESSEL MORPHOGENESIS
CIRCULATORY SYSTEM DEVELOPMENT
TUBE MORPHOGENESIS
Up−regulated
ANTIGEN PROCESSING AND PRESENTATION OF PEPTIDE OR POLYSACCHARIDE ANTIGEN VIA MHC CLASS II
ANTIGEN PROCESSING AND PRESENTATION OF PEPTIDE ANTIGEN
DETOXIFICATION
ANTIGEN PROCESSING AND PRESENTATION OF EXOGENOUS PEPTIDE ANTIGEN VIA MHC CLASS II
CELLULAR OXIDANT DETOXIFICATION
PROTEIN FOLDING
MONOCARBOXYLIC ACID METABOLIC PROCESS
CARBOHYDRATE DERIVATIVE METABOLIC PROCESS
ORGANIC ACID METABOLIC PROCESS
SMALL MOLECULE METABOLIC PROCESS
NES
M
Cluster 13 - Mesothelial cells
Down−regulated
SYNAPSE ASSEMBLY
NEURON RECOGNITION
NEUROTRANSMITTER SECRETION
EXTERNAL ENCAPSULATING STRUCTURE ORGANIZATION
SYNAPSE ORGANIZATION
DENDRITE DEVELOPMENT
CELL JUNCTION ORGANIZATION
NEURON DEVELOPMENT
CELLULAR COMPONENT MORPHOGENESIS
CELL CELL SIGNALING
Up−regulated
CHAPERONE COFACTOR DEPENDENT PROTEIN REFOLDING
DE NOVO PROTEIN FOLDING
CHAPERONE MEDIATED PROTEIN FOLDING
PROTEIN TARGETING TO MITOCHONDRION
MITOCHONDRIAL TRANSPORT
MITOCHONDRION ORGANIZATION
REGULATION OF APOPTOTIC SIGNALING PATHWAY
ATP METABOLIC PROCESS
GENERATION OF PRECURSOR METABOLITES AND ENERGY
APOPTOTIC SIGNALING PATHWAY
REGULATION OF CELL DEATH
NES
N
Cluster 15 - Red blood cells (RBCs)
Down−regulated
CELLULAR RESPONSE TO CALCIUM ION
CELLULAR RESPONSE TO ANTIBIOTIC
MUSCLE STRUCTURE DEVELOPMENT
NES
Up−regulated
No enrichment
GOBP POSITIVE REGULATION OF INFLAMMATORY RESPONSE
O
Estrus_INFAMP
P
Aged_INFAMP
umap_2
umap_1

**Suppl. Fig. 5: GSEA of each cluster showing genesets down- or upregulated in 18mo INF/AMP cells, relative to 3mo INF/AMP cells.** (A-C) GSEA showing down- and upregulated genesets in SC clusters 1 (A), 2 (B), and 11 (C). (D-F) GSEA showing down- and upregulated genesets in stromal cell clusters 3 (D), 7 (E), and 10 (F). (G-I) GSEA showing down- and upregulated genesets in muscle cell clusters 5 (G), 9 (H), and 14 (I). (J, K) GSEA showing down- and upregulated genesets in immune cell clusters, including cluster 12, identified as T-cells (J), and cluster 8, identified as antigen presenting cells/APCs (K). (L) GSEA showing down- and upregulated genesets in cluster 6, identified as endothelial cells. (M) GSEA showing down- and upregulated genesets in cluster 13, identified as mesothelial cells. (N) GSEA showing down- and upregulated genesets in cluster 15, identified as red blood cells/RBCs. (O, P) UMAP showing expression of gene set associated with positive regulation of inflammatory response in young (O) and aged (P) INF/AMP cells.

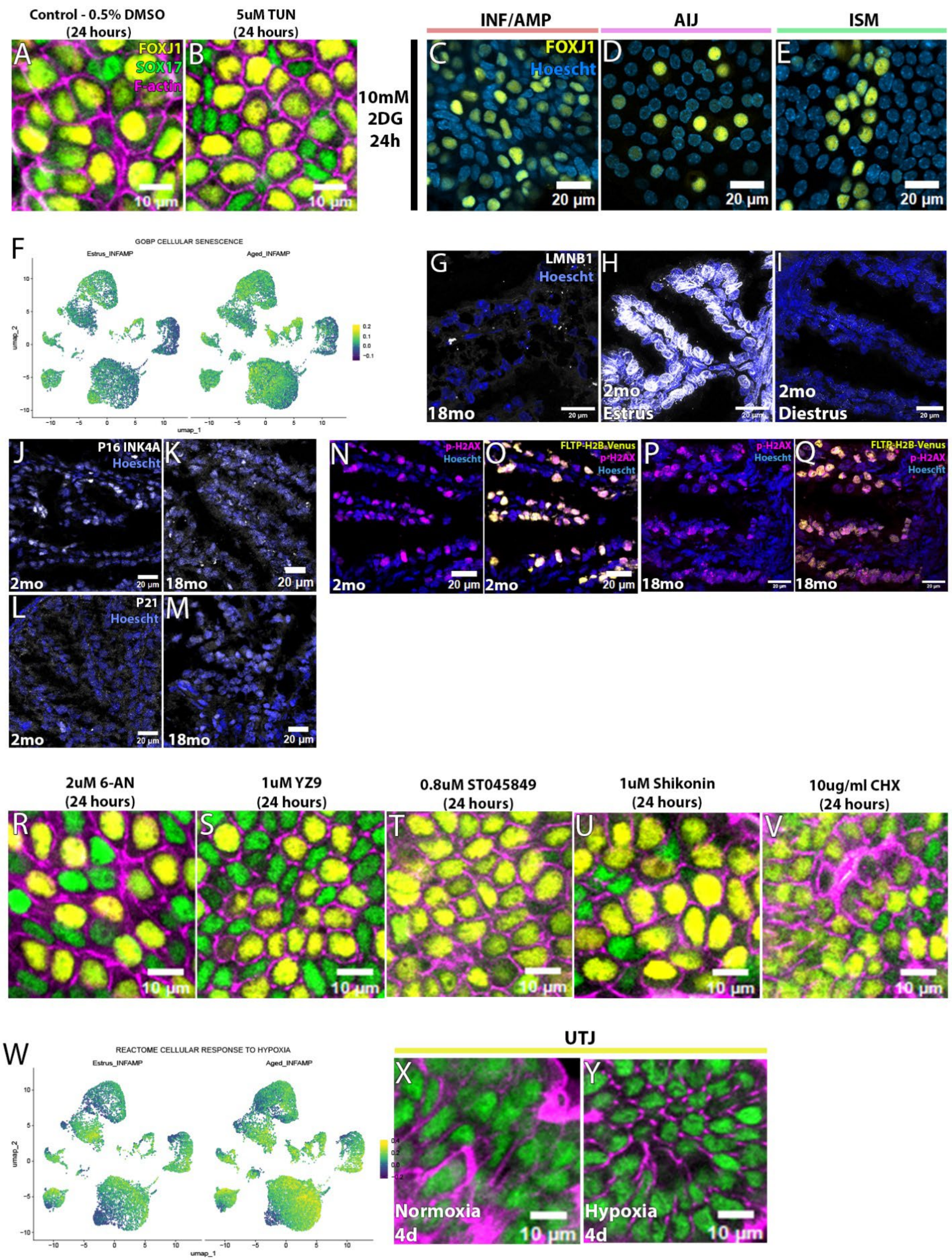
Control - 0.5% DMSO (24 hours)
5uM TUN (24 hours)
FOXJ1
SOX17
F-actin
10 µm
INF/AMP
AIJ
ISM
10mM 2DG 24h
FOXJ1
Hoescht
20 µm
GOBP CELLULAR SENESCENCE
Estrus_INFAMP
Aged_INFAMP
umap_1
umap_2
LMNB1
Hoescht
18mo
2mo Estrus
2mo Diestrus
20 µm
P16 INK4A
Hoescht
2mo
18mo
P21
Hoescht
p-H2AX
Hoescht
FLTP-H2B-Venus
p-H2AX
Hoescht
2uM 6-AN (24 hours)
1uM YZ9 (24 hours)
0.8uM ST045849 (24 hours)
1uM Shikonin (24 hours)
10ug/ml CHX (24 hours)
REACTOME CELLULAR RESPONSE TO HYPOXIA
Estrus_INFAMP
Aged_INFAMP
UTJ
Normoxia 4d
Hypoxia 4d
10 µm

**Suppl. Fig. 6: MCC vacuolation is unaffected by ER stress induction, or inhibition of glycolysis, pentose phosphate, or hexosamine pathways.** (A, B) ER stress was induced by treating with Tunicamycin/TUN for 24 hours in INF/AMP organotypic slice cultures. No obvious cellular differences were noted in treated cultures (B) and controls (A). (C-E) Inhibition of glycolysis in organotypic slice cultures. 24-hour-long 2DG treatment did not induce any obvious cellular differences in the INF/AMP (C), AIJ (D), or ISM (E). (F-K) Cellular senescence markers in the INF/AMP region of aged and young mice. Expression of cellular senescence-associated geneset between young (F) and aged (G) INF/AMP cells. Distinct LaminB1 expression levels in the INF/AMP region of 2mo mice in estrus (H) and diestrus stages (I). Lamin B1 was not detected in 18mo INF/AMP (J). Similar expression patterns of phospho-H2AX (K, L), P16 (M, N), and P21 (O, P) in young and aged mice. (P-T) Induction of metabolic stress/nutrient deprivation in INF/AMP slice cultures. No cellular differences were noted upon inhibition of pentose phosphate pathway (P), hexosamine biosynthesis pathway (Q), PFKFB3 (R), PKM2 (S), and protein synthesis (T). (U, V) UMAP showing expression of geneset associated with cellular response to hypoxia in young (U) and aged (V) INF/AMP cells. (W, X) No cellular changes in UTJ epithelium in normoxia (W) and hypoxia (X).

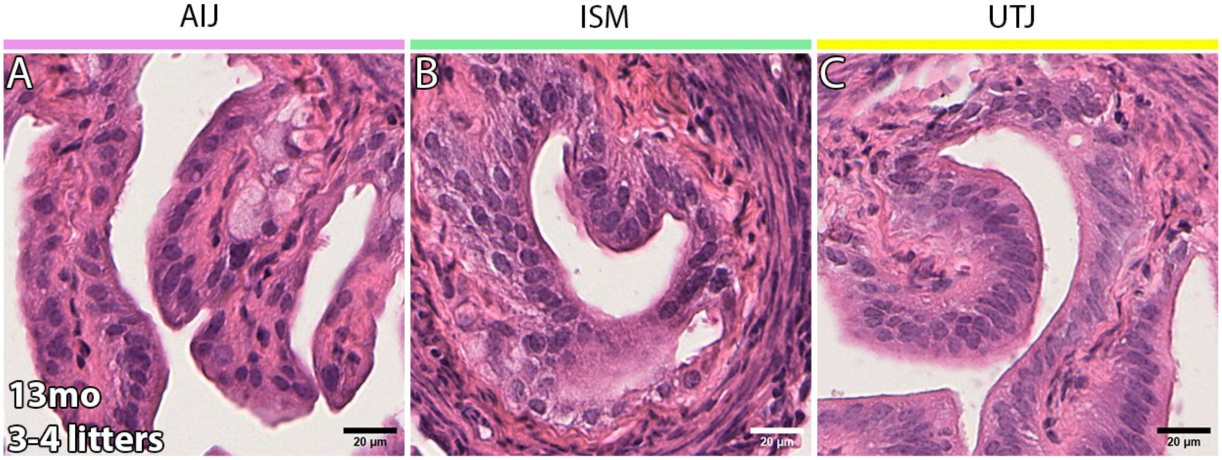

**Suppl. Fig. 7: No discernible vacuoles in the AIJ, ISM, and UTJ regions of 13mo mice with 3-4 litters.** (A-C) Vacuoles were not reproducibly observed in the AIJ (A), ISM (B) and UTJ (C) regions of 13mo mice that had given birth to 3-4 litters.

**Suppl. Vid. 1: Cilia beating continued regardless of aging.** (A-C) Live imaging of cilia beating in 2mo (A) and 13mo (B) INF/AMP.
